# Joint inference of adaptive and demographic history from temporal population genomic data

**DOI:** 10.1101/2021.03.12.435133

**Authors:** Vitor A. C. Pavinato, Stéphane De Mita, Jean-Michel Marin, Miguel de Navascués

## Abstract

Disentangling the effects of selection and drift is a long-standing problem in population genetics. Simulations show that pervasive selection may bias the inference of demography. Ideally, models for the inference of demography and selection should account for the interaction between these two forces. With simulation-based likelihood-free methods such as Approximate Bayesian Computation (ABC), demography and selection parameters can be jointly estimated. We propose to use the ABC-Random Forests framework to jointly infer demographic and selection parameters from temporal population genomic data (e.g. experimental evolution, monitored populations, ancient DNA). Our framework allowed the separation of demography (census size, *N*) from the genetic drift (effective population size, *N*_e_) and the estimation of genome-wide parameters of selection. Selection parameters informed us about the adaptive potential of a population (the scaled mutation rate of beneficial mutations, *θ*_b_), the realized adaptation (the number of mutation under strong selection), and population fitness (genetic load). We applied this approach to a dataset of feral populations of honey bees (*Apis mellifera*) collected in California, and we estimated parameters consistent with the biology and the recent history of this species.

## Introduction

One aim of population genomics is to understand how demography and natural selection shape the genetic diversity of populations. A classical approach assumes that demography (migration, population subdivision, population size changes) leaves a genome-wide signal. In contrast, selection leaves a localized signal close to where the causal mutation is located. Many methods follow this approach to infer demography or selection (reviewed by Beichman et al., 2018; Casillas and Barbadilla, 2017). Demographic inference assumes that most of the genome evolves without the influence of selection and that any deviation from the mutation-drift equilibrium observed in the data was caused by demographic events (Beichman et al., 2018). Many of the methods search for locus-specific signals of selection left on nearby neutral mutations (Fay and Wu, 2000; Kim and Nielsen, 2004; Tajima, 1989) (low genetic diversity and high differentiation) to localize the region affected by selection mutation, assuming a specific demography (constant population size in early methods; Nielsen, 2005; Pool et al., 2010).

Conducting demographic and selection inference separately may have some shortcomings. First, there is the assumption that the signal left by demography is little affected by selection because selection is rare. However, linked selection can affect neutral and weakly selected sites that are far from the mutation targeted by selection (Neher, 2013; Sella et al., 2009) and selection can be pervasive (Lange and Pool, 2018; Sella et al., 2009). In addition, some methods for selection scans are not robust to misspecifications of demographic history. Consequently, an unspecified bottleneck or population increase, for example, can inflate the false positive rate of genome scans (Jensen, Kim, et al., 2005; Jensen, Thornton, et al., 2007; Schrider, Shanku, et al., 2016). These findings highlight the necessity of inferential methods that jointly accounts for the multiple evolutionary forces that act on populations (Bank et al., 2014; J Li et al., 2012; Lin et al., 2011).

Some methods to perform the Joint inference of demography and selection have been proposed by making the explicit assumption that a set of neutral polymorphisms can be distinguished from another set of polymorphisms putatively under selection (usually synonymous *vs*. non-synonymous sites). McDonald and Kreitman (1991) first proposed to compare polymorphism and divergence between synonymous and non-synonymous sites and soon after Sawyer and Hartl (1992) proposed to model the evolution both types of sites with a Poisson random field. Several developments based on these ideas have been put forward over the years, improving and refining the models (*e.g*. Boyko et al., 2008; Messer and Petrov, 2013). Alternatively, other approaches do not make the assumption about the neutral nature of some specific polymorphisms. In these, the effect of selection is assumed to be heterogeneous among loci or genomic regions, and they often have the additional objective to identify the loci or regions under (strong) selection. In the models used by these approaches, it is often difficult to calculate the likelihood (but see Vitalis et al., 2014). Methods that rely on simulations provide easier alternatives to using likelihood functions (Csilléry et al., 2010; Schrider and Kern, 2018). One of the first works that proposed such a strategy addressed the inference of local adaptation (Bazin et al., 2010). With coalescent simulations of an island model, Bazin et al. (2010) estimated demographic parameters and inferred the number of loci under selection. In their simulations, the selection was modeled as locus-specific migration rates in which a selected locus had lower migration rates than neutral loci. However, locus-specific migration rates or effective population size (as in Fraïsse et al., 2021; Roux et al., 2016) represent crude approximations of the selection process. Forward-in-time simulation allows more realistic models of selection. These were used to make inferences on *N*_e_ in the presence of selection by Sheehan and Song (2016) (selective sweeps and balancing selection) and Johri et al. (2020) (background selection). This strategy brought new insights into the dynamics of selection. Laval et al. (2019) estimated the number of past selective sweeps in the human genome in the past 100,000 years, their intensity, and their age. These works exemplify the power of likelihood-free methods to infer the complex interaction between demography and selection. However, all the methods discussed above rely on independent simulation or modeling of loci or genomic regions which prevents the modeling of genome-wide effects of selection as the reduction of effective population size due to the variance of reproductive success of individuals (Santiago and Caballero, 1995) or the combined effects of mutations on individual fitness.

Most population genetic studies use samples collected at a one-time point to infer the neutral processes (mutation, recombination, random genetic drift) and selection throughout the history of populations. Temporal data allows a better understanding of recent evolutionary processes (e.g. Dehasque et al., 2020; Feder, PS Pennings, et al., 2021) because they contain information about the allele frequency changes through time. By tracking the allele frequency changes overtime, it is possible to estimate the relative role of selection and drift. Consequently, temporal data has the potential to give us a better understanding of the interaction between drift and selection (see for example, Buffalo and Coop, 2019, 2020).

Here, we propose using ABC to jointly estimate demography and positive selection from temporal genomic data. In our framework, we use individual-based, forward-in-time simulations, which allow the modeling of the genome-wide, linked selection and additive effects of beneficial mutations. Until recently, such computationally demanding simulations in ABC inference were unrealistic since many simulations are required to achieve accuracy in ABC (Frazier et al., 2018). However, with the introduction of Random Forests (ABC-RF), it is now possible to reduce the computational burden as fewer simulations are required to achieve reliable estimates (Pudlo et al., 2016; Raynal et al., 2019). While many methods focus on the detection of targets of selection, our work addresses the inference of parameters that characterizes the genome-wide signal of demography and selection. Our genome-wide estimates were reasonably accurate for a wide range of adaptation rates and strength of selection. We were able to separate the estimates of *N*_e_ (a measure of genetic drift) from the population census size *N*. We also estimated the influx of new beneficial mutations as measured by the population scaled mutation rate of beneficial mutations. The separation between demography and drift and the inference of genome-wide selection was only possible using latent variables. Latent variables emerged as properties of each simulation, and consequently, they better captured the emerging interaction between demography and selection than model parameters. We first evaluated the performance of an ABC-RF approach with forward-in-time simulations. Finally, we applied this framework to the analysis of a real time-series population genomics dataset of the feral population of honey bees (*Apis mellifera*, Cridland et al., 2018). Our results were consistent with the species’ biology and with events that occurred recently in the history of the analyzed populations, taking into account the limitations of the current implementation of our approach.

## Material and methods

### Inference model

We assumed a closed population (no migration) of *N* diploid individuals that evolved under a Wright-Fisher model with selection. The population census size *N* was constant, and selection only acted on *de novo* beneficial mutations that were allowed to arise in the population since the first generation (generation one corresponds to the first burn-in generation). Every beneficial mutation had a selection coefficient of *s* higher than zero, and all were co-dominant. The values of the selection coefficients s were drawn from a gamma distribution with mean *γ* and scale parameter 1. Beneficial mutations entered the population with a rate of *μ*_b_ per generation independent of the mutation selective strength. Consequently, we defined the scaled mutation rate of the beneficial mutations per generation *θ*_b_ as the product the population size *N*, the mutation rate of beneficial mutation *μ*_b_ and the genome size *G*, *θ_b_* = 4*Nμ*_b_*G*. This rate determines the amount of new beneficial mutations that arise in the population every generation. It can also be viewed as the waiting time for the appearance of a new beneficial mutation in the population. Populations with high *θ*_b_ receive new beneficial mutations every generation (Karasov et al., 2010), but a population with low *θ*_b_ needs to wait more time for a new beneficial mutation to arise.

We divided the model into two periods: 1)the burn-in period, which is necessary to remove from the simulations anyfootprint of the initial simulation state; the duration of this period was defined as the time necessary to reach a point were the most recent common ancestors for all genomic regions are more recent than the start of the simulation (*i.e*. the burn-in was run until this condition was fulfilled); and 2) the inference period, where we defined the longitudinal samples of individuals. These two periods were defined by their time spam and the population census size, being *N*_0_ and *N* as the population size of the burn-in and the inference period, respectively. Population size is constant within each simulated period and changes between periods.

First sample of individuals was taken at *t*_1_, the immediate next generation after burn-in ended; the second was taken at *t*_2_, after *τ* generations from *t*_1_. Individuals were sampled following the sample plan II of Nei and Tajima (1981), where individuals were taken before reproduction and permanently removed from the population. In this way, their genotypes did not contribute to the next generation.

Each individual’s genome of size *G* (in base pairs) consisted of a single linkage group with a per base recombination rate per generation of *r*. We modeled the selection effect in this genome by dividing it into “neutral” and “non-neutral” regions. Non-neutral regions held both neutral and beneficial mutations. This division can be interpreted as a genomic architecture in which genic regions have a combination of neutral (synonymous intron mutations) and selected (non-synonymous mutation) sites with intergenic regions (neutral mutations) in between. However, this architecture allowed simulating the heterogeneous selection action along the genome.

We chose this simplification because it is a general and straightforward way to define independent priors for the relative number of non-neutral to neutral regions and for the number of beneficial mutations in nonneutral regions. The probability of beneficial mutation to arise in the simulation (*i.e*. the mutation rate per generation, *μ*_b_) was determined by the product of the proportion of non-neutral regions *P*_R_, the proportion of beneficial mutation in a non-neutral region *P*_B_ and the mutation rate per generation *μ*. Figure 1 shows a schematic representation of the model template (and see Table SI for a summary of the notation).

**Figure 1.**
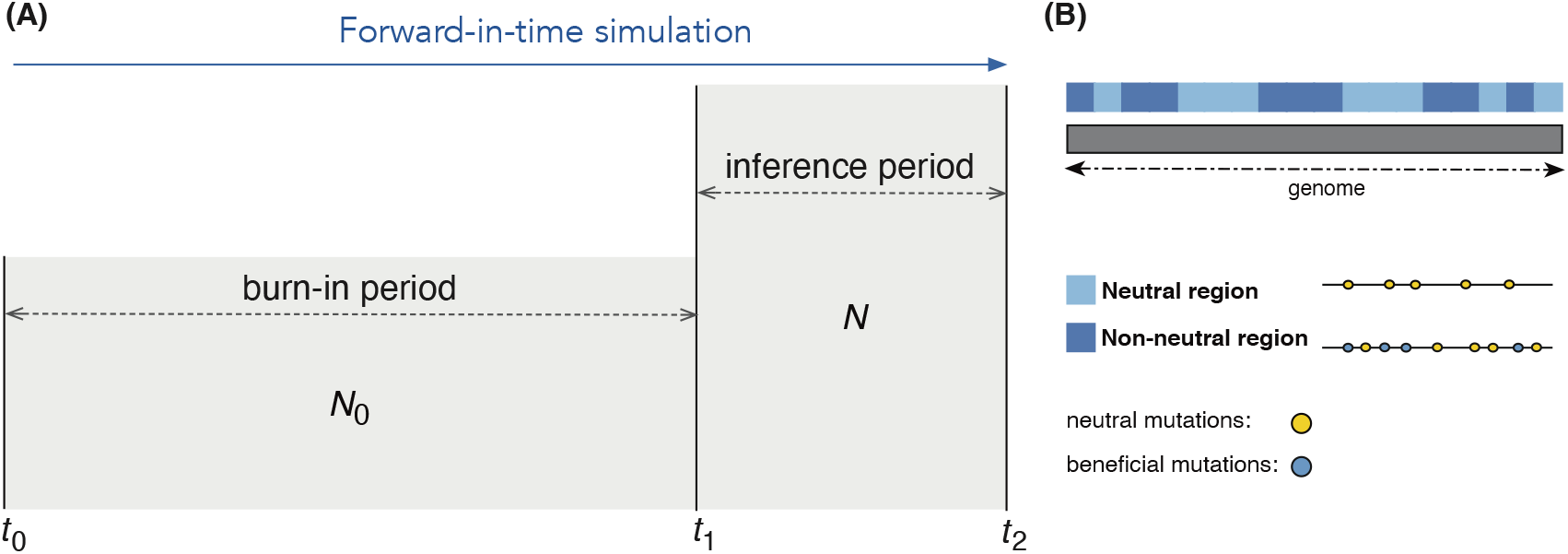
A schematic representation of the model used to simulate temporal population genomic data. (A) the population model consisted of 1) the burn-in period, where the numberof generations was determined by the time necessary to contain the MRCA for all genomic regions. 2) the inference period between the two time points, where the inference of demography and selection was made. (B) the genomic architecture model consisted of 1) a diploid genome of one linkage group divided into neutral and non-neutral regions composed of neutral and a combination of neutral and beneficial mutations. Note that, despite the graphical representation, the model does not condition *N* to be larger than *N*_0_, both expansions and contraction are considered in the model.

### Calculation of summary statistics and latent variables

The above model was used to simulate the dynamic of drift and selection in a closed population. In the two sample periods, individuals from the whole population were sampled and used for the calculation of the summary statistics for the ABC-RF framework. For each simulation, we calculated summary statistics that: 1) compared the two samples (*e.g*. genetic differentiation, *F*_ST_), and 2) quantified the diversity within-sample (*e.g*. expected heterozygosity, *H*_E_). For the latter, statistics were obtained for each and all pooled samples. Some summary statistics were calculated genome-wide. For example, global *F*_ST_, global *H*_E_ and the total number of polymorphic sites *S*; others were calculated SNP-by-SNP as the *H*_E_; or they were calculated in windows as *S*, the nucleotide diversity *π*, and Tajima’s *D*. For every simulation, we measured the mean, variance, kurtosis, skewness, and 5% and 95% quantiles among all locus-specific or window summary statistics. These statistics inform about the heterogeneity of genome-wide distribution of locus-specific or window summary statistics. We set three window sizes for the window summary statistics: 500,5,000, and 10,000 bp. Windows overlapped because each was composed around every SNP, putting the targeted variation in the middle of the window with other surrounding SNPs in half the window size on each side of the targeted SNP. The site-frequency spectrum was obtained as a global summary statistics with three different numbers of discrete classes (bin sizes): 10, 15, and 20 bins (the complete list of summary statistics can be found in Supplementary Methods, section S1.1 List of summary statistics).

For every simulation, we combined a vector of summary statistics with the vector of X model parameters and the vector of five latent variables. Latent variables represent values from the simulation or that emerged by combining a latent variable and a model parameter. In our inferential framework, for example, the effective population size *N*_e_ is a latent variable calculated within each simulation. The ratio between the effective population size *N*_e_ and the population census size *N*, *N*_e_/*N*, on the other hand, was derived by combining a latent variable and a model parameter for each simulation. The other three latent variables were: the number of beneficial mutations under strong selection *P*, the average selection coefficient of strongly selected mutations 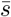, and the average substitution load *L*.

The effective population size *N*_e_ measures the increase of inbreeding at each generation. In this definition, *N*_e_ is the size of an ideal population with the same amount of drift as the population under consideration. Defined in these terms, *N*_e_ is the inbreeding effective size (Santiago and Caballero, 1995; Walsh and Lynch, 2018). It was calculated in every generation *i* of the sampling period as:

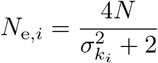

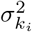 being the variance among parents of the number of gametes produced that contributed to offspring in generation *i*. The *N*_e_ for the whole inference period was obtained by calculating the harmonic mean of *N*_e,*i*_. The population size of *N* was kept constant for the whole period, as shown above, representing a simulation parameter. From the *N*_e_ we obtained the ratio *N*_e_/*N* (it measures how the census size reflects the actual effective population size: we expect to have a reduction on *N*_e_ compared to *N* when beneficial mutations are more pervasive).

We also recorded the selection coefficient of all beneficial mutations present in every generation *i* from *t*_1_ to *t*_2_ in each simulation. After, we calculated the fraction of beneficial mutations that were strongly selected (where *s* > 1/*N*_e_ over all mutations that were segregating in the period). This fraction represented all beneficial mutations present between *t*_1_ and *t*_2_, regardless if they were lost or fixed at any generation of the period or if their frequency fluctuated but never reached fixation. We decided on it because any beneficial mutation can impact the allele frequency trajectories of other mutations (neutral or beneficial). For these mutations, we also calculated the average across all selection coefficients. We also calculated, in every generation of this period, the substitution load *L_i_* as the difference between the total fitness of the individual with the highest fitness *W*_max*i*_ and the mean total fitness of the population 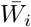 (it measures the overall diversity of beneficial mutations present in the inference period),

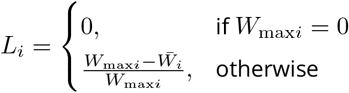

The average substitution load was obtained by averaging all values of *L_i_*.

### Implementation

The model was simulated with the software SLiM v3.1 (Haller, Galloway, et al., 2019; Haller and Messer, 2017). To calculate the inbreeding effective size, we needed to activate an optional SLiM 3.1 behavior to track the pedigrees of each individual in the population. It allowed us to obtain the number of each parent gamete and the population variance of the number of gametes. For calculating the generation substitution load, we used a SLiM built-in function that allowed us to obtain the fitness vector of all individuals in the population. The cached fitness was the sum of all fitness determined by each beneficial mutation.

Each simulation was produced by using different combinations of the model’s parameters: 1) the mutation rate per bp per generation *μ*, 2) the per-base recombination rate per generation *r*, 3) the mean *γ* of a gamma distribution (with the shape parameter equal to the mean), from which the selection coefficients *s* of each beneficial mutation in the simulation were sampled, 4) the number of non-neutral genomic regions *P*_R_, 5) the parameter that determines the probability of beneficial mutation in non-neutral regions *P*_B_, 6) the population census size of the burn-in period *N*_0_, and, finally, 7) the population size of the inferential period *N*.

We set SLiM to output genotypic data of samples of individuals as single nucleotide polymorphisms (SNPs), at *t*_1_ and *t*_2_, in the VCF file format. Using bcftools (H Li, 2011), custom R function (R Core Team, 2020) and EggLib (Siol et al., 2022), SLiM outputs were processed and summary statistics calculated. We implemented a pipeline in an R script that automates the sampling of the prior values, runs each simulation, manipulates the VCF files, calculates the summary statistics, and organizes the final reference table. This script was also produced to facilitate the model test with a few simulations and the job submission in an HPC node(s). The main R and additional scripts are available on Zenodo (Pavinato et al., 2022). In this pipeline, for every simulation, a row of the reference table was produced by combining the model parameters, latent variables, and summary statistics.

### ABC-RF

In this work, we use Random Forests (RF) in the ABC procedure, where the parameter estimation is a machine learning problem (Pudlo et al., 2016; Raynal et al., 2019). The performance of this approach was evaluated through simulations. First, we assumed a target dataset consisting of two samples of 100 individuals sampled ten generations apart from the same population. A reference table for that target data was produced by simulating the whole-genome SNPs of diploid individuals using the model described above and calculating the previous summary statistics. At each simulation, we sampled 100 individuals at each time point and recorded their genotypes. Only polymorphic SNPs were retained for each sample. In each simulation, each individual had a genome size of 100 Mbp divided into 2,000 fragments of 50,000 bps. A number of these fragments were randomly set as either neutral or non-neutral, based on the probability *P*_R_. For all model parameters, values of each simulation were sampled from a log-uniform distribution with range: 1 to 2,000 for *N*_0_ and *N*, 10^-10^ to 10^-6^ for *μ*, 5 × 10^-10^ to 5 × 10^-7^ for *r*, 10^-5^ to 1 for *P*_B_, and 10^-3^ to 1 for *γ*. Furthermore, uniform distribution with range 0 to 1 for *P*_R_ (Figure SI shows the prior distribution for all model parameters and latent values).

The raw reference table produced by the pipeline was processed to remove missing data. Missing data were present in several summary statistics of simulations with low genetic diversity that can be produced, for example, by low mutation rate, small population size, selection, or the combination of these parameters. Missing data were also present in the entire row of a simulation if the combination of population size, mutation, and especially recombination rate produced simulations that were memory intense, which caused the simulation to crash. A final reference table containing 55,634 simulations with 405 summary statistics was used to train the ABC-RFs. Independent RFs were obtained for each parameter and latent variable using R package `abcrf` (Pudlo et al., 2016; Raynal et al., 2019). Each RF was obtained by growing 1,000 trees. The RFs were grown with the default parameters. Average genetic load, *L*, and *P* were logit-transformed before the training. For these latent variables and for 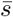, simulations with *L* = 0, *P* = 0 or 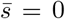 were also excluded from the training set, which reduced it to 36,026 simulations for *L*, and with 29,264 simulations for *P* and 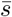. We performed log transformation before training for the other parameters and latent variables and used the reference table containing all simulations.

The performance of each trained Random Forest was evaluated with *out-of-bag* (OOB) estimates (Breiman, 2001). The trained model produced these estimates for the data used for training. Regression trees that compose the actual RF are grown using part of the data selected randomly from the initial set of simulations. Consequently, for each simulation, a subset of trees was grown without the data from that simulation. The estimate from that subset of trees is called the OOB estimate, and with it, the trained model is validated without splitting the reference table into the training and testing sets. We calculated the mean squared error (MSE) and the correlation coefficient (*R*^2^) between the true and the OOB estimated values obtained with the function `regAbcrf` implemented in the R package `abcrf`. For neutral simulations of the latent variables *L*, *P*, and 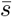, we evaluated the performance with the MSE and the bias on the parameters estimated in the original parameter scale.

An additional 1,000 simulations were used to evaluate the method’s robustness to heterogeneous recombination rates along the genome. The simulation model was identical to the previously described simulations, except that a recombination map was used with varying recombination rates along the genome. We used the already implemented genomic fragmentation of the genome in “neutral” and “non-neutral” regions, which split the genome into 2,000 blocks of 50 Kbp, to define the positions at which the recombination rate changed. Each corresponding fragment had a recombination rate sampled from a log-uniform distribution with a range between 10^log_10_ *r*-0.5^ and 10^log_10_ *r*+0.5^, with *r* sampled from the prior distribution as described above. This range allowed the genome to have recombination rates spanning one order of magnitude. We evaluate the RF performance in these simulations by calculating the mean squared error (MSE) and the correlation coefficient (*R*^2^) between the true parameter values and the RF estimates. For neutral simulations of the latent variables *L*, *P*, and 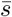, we evaluated the performance with the MSE and the bias.

### Alternative estimates of *N*_e_ from temporal data

We compared the ABC-RF *N*_e_ estimates with estimates obtained with the global *F*_ST_ between temporal genomic samples (Frachon et al., 2017). This estimator is defined as:

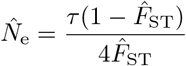

where *τ* accounts for the time-interval, in generations, between the first and the last samples used to estimate the *F*_ST_, and 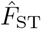 is the Weir and Cockerham’s *F*_ST_ estimator (Weir and Cockerham, 1984). The *N*_e_ from the *F*_ST_ was calculated for all simulations used to train the random forest. We calculated the mean squared error (MSE) and the squared correlation coefficient of linear regression (*R*^2^) between the observed (true) and the *F*_ST_-based *N*_e_ estimated values of all simulations. We also evaluated the performance of each estimator by calculating the MSE for simulations within a specific range of values of *θ*_b_ (local MSE estimates). By comparing the changes in MSE values of each estimator as a function of *θ*_b_ we could better understand how the amount of selection affected each estimator.

### Analysis of temporal genomic data of feral populations of *Apis mellifera*

We used our framework to analyze the whole-genome sequencing data of feral populations of honey bees from California (Cridland et al., 2018). Eight out of fourteen sites in this work were composed of samples from museum and contemporary collections of freely foraging honey bees: 1) Avalon site in Catalina Island, Los Angeles County, 2) Arcata and Blue Lake sites in Humboldt county, 3) Placerita Canion Nature Area in Los Angeles County, 4) Sky Valley and Idyllwild in Riverside County, 5) La Grange, Stanislaus county, 6) Stebbins Cold Canyon Reserve, Solano county and 7) UC Davis Campus, Yolo county (Table 1). This dataset contains pairs spanning 104 years (as in the Avalon site, Catalina Island, Los Angeles county) and pairs spanning only 15 years (as in the Placerita Canyon Nature Area, Southern California, and Idyllwild, in Riverside county). For the temporal samples from Riverside County, we only used the two samples collected in May 1999 in Idyllwild as the first sample. We combined all samples collected in September 2014 (in Idyllwild and Sky Valley) as the second sample (Table 1). Publicly available whole genomes fastq files for the contemporary and museum samples are available from the Sequence Read Archive (PRJNA385500) as described by Cridland et al. (2018); we performed the data analysis from VCF files (the same files used in Cridland et al., 2018) available in Pavinato et al. (2022).

**Table 1.**
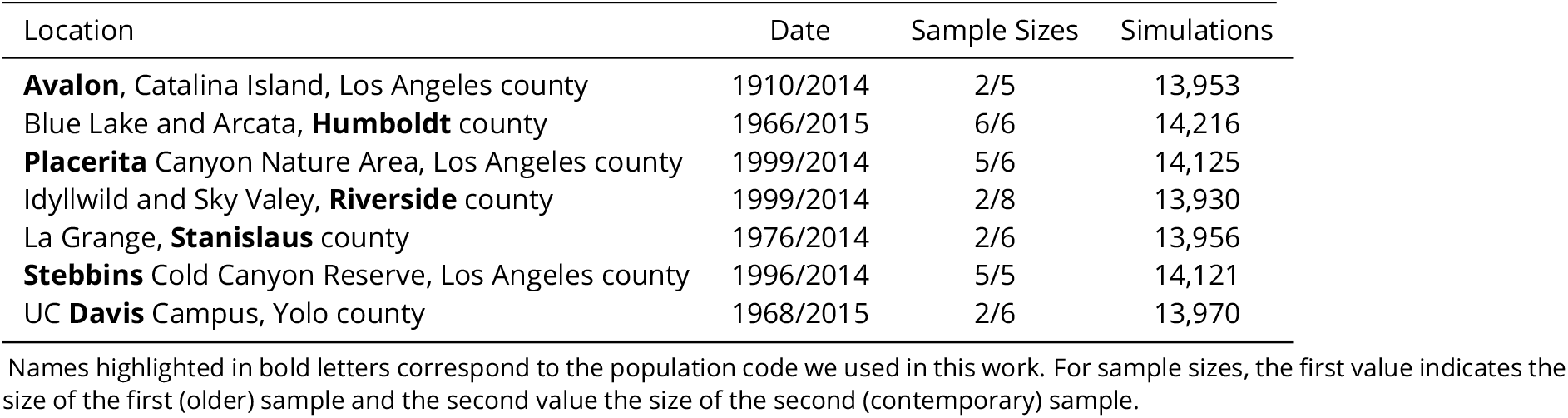
Populations and number of simulations in the reference table.

| Location | Date | Sample Sizes | Simulations |
| --- | --- | --- | --- |
| **Avalon**, Catalina Island, Los Angeles county | 1910/2014 | 2/5 | 13,953 |
| Blue Lake and Arcata, **Humboldt** county | 1966/2015 | 6/6 | 14,216 |
| **Placerita** Canyon Nature Area, Los Angeles county | 1999/2014 | 5/6 | 14,125 |
| Idyllwild and Sky Valey, **Riverside** county | 1999/2014 | 2/8 | 13,930 |
| La Grange, **Stanislaus** county | 1976/2014 | 2/6 | 13,956 |
| **Stebbins** Cold Canyon Reserve, Los Angeles county | 1996/2014 | 5/5 | 14,121 |
| UC **Davis** Campus, Yolo county | 1968/2015 | 2/6 | 13,970 |

Names highlighted in bold letters correspond to the population code we used in this work. For sample sizes, the first value indicates the size of the first (older) sample and the second value the size of the second (contemporary) sample.

Individual VCF files of each population were combined with bcftools (H Li, 2011), and a custom R script was used to convert each dataset to the input format required to run an EggLib custom implementation (in Pavinato et al., 2022). We first produce simulated data to train the RF to apply our model to this targeted dataset. A reference table was produced by simulating whole-genome SNPs for diploid individuals of *Apis mellifera*, changing three model parameters specifically for this targeted dataset: the sample size for population time points *t*_1_ and *t*_2_, and the size of the haploid genome. For each population, we set the simulation to sample the same number of sequenced individuals from the pool of simulated individuals (as detailed in Table 1). For the Avalon population, for example, at *t*_1_ and *t*_2_ we set the simulation to sample two and five individuals apart *τ* = 104 generations (assuming one generation/year). Only polymorphic SNPs were retained for each sample. We set the haploid genome size to 250 Mbp (similar to the most recent estimates of *A. mellifera* genome size; Elsik et al., 2014). We measured the amount of missing data present in the original VCF files (Cridland et al., 2018) for each population. We found a negligible amount (< 1%) in most of the populations (except populations from Avalon and Placerita that had 10% of the total missing genotypes), and we decided not to simulate missing data for any population analyzed.

The simulation model to generate the reference tables for the ABC analysis of *A. mellifera* populations was similar to the model described above but required some modification to adjust it to the specificities of the species and samples available. The simulated genome was divided into 5,000 fragments of 50,000 bps. These fragments were randomly set as neutral or non-neutral according to the parameter *P*_R_. We used a Normal distribution for *μ* with a mean of 3.4 × 10^-9^ with a standard deviation of 0.5 to have a prior distribution center around the estimated mutation rate for Hymenoptera (Liu et al., 2017). The per base recombination rate was set as Uniform, ranging from 10^-8^ to 10^-4^. A single linkage group represented the genome. The population sizes *N*_0_ and *N* were taken from a Uniform prior distribution ranging from 1 to 10,000 individuals. Other prior probability distributions of the parameters were set with the same prior as described above. Sample sizes and times were adjusted to match each population’s population (see table 1). We used the same summary statistics described above. However, we calculated only one window size of 10Kbp for summary statistics calculated in windows and one bin size of 10 bins for the site-frequency spectrum. The raw reference table containing the vector of parameters, latent variables, and summary statistics produced by the pipeline was processed to remove missing data. A final reference table containing 162 summary statistics for each population pair was used to train the ABC-RFs. We visually assessed the model goodness-of-fit by performing a principal component analysis on the summary statistics of each population training reference table and projecting the corresponding PC of the target population reference table on the PCA plot. We consider a good model fit when the target population data point falls within the cloud of population simulated data points.

Like the ABC analyses described above, independent RFs were obtained for each parameter and latent variable using R package `abcrf` (Pudlo et al., 2016; Raynal et al., 2019). Each RF was obtained by growing 1,000 trees. The RFs were grown with the default parameters. Average genetic load, *L*, and *P* were logit-transformed before the training. For these latent variables and for 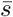, simulations with *L* = 0, *P* = 0 or 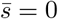 were also excluded from the training set. We performed log transformation before training for the other parameters and latent variables and used the reference table containing all simulations. As before, we evaluate the RF performance by calculating the mean squared error (MSE) and the correlation coefficient (*R*^2^) between the true and the OOB estimated values obtained with the function `regAbcrf` implemented in the R package `abcrf`. For neutral simulations of the latent variables *L*, *P*, and 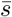, we evaluated the performance with the MSE and the bias on the parameters estimated in the original parameter scale. See Table 1 for the number of simulations of each reference table.

## Results

### Joint inference of adaptive and demographic history

The proposed framework allowed us to estimate parameters informative about adaptive and demographic history in temporal population genomics settings. Independent random forests estimated the population scaled beneficial mutation rate *θ*_b_, the population census size *N*, and the effective population size *N*_e_ (Figure 2). Trained RFs performed well in predicting *N* and *N*_e_ with small MSE and higher *R*^2^ (Figure 2 b and c). But, the trained RF for *θ*_b_ had a lower performance than the trained RFs for demographic parameters, with high MSE and low *R*^2^ (Figure 2a). Still, the estimates were robust for intermediate to higher values of *θ*_b_. For the results of other model parameters and latent variables informative about demography and selection (see Figure S2 and section S2 Supplementary Results). Similar values of MSE and *R*^2^ on true vs. RF estimated values (Figure S5 a, b, and c) indicated similar performance RF for *θ*_b_, *N*, and *N*_e_ on simulations with heterogenous recombination rates (see Figure S4 for an example of how *r* could vary across the genome). For the results of other model parameters and latent variables for simulation with heterogenous recombination rate, see Figure S5 and Table S2, section S2 Supplementary Results.

**Figure 2.**
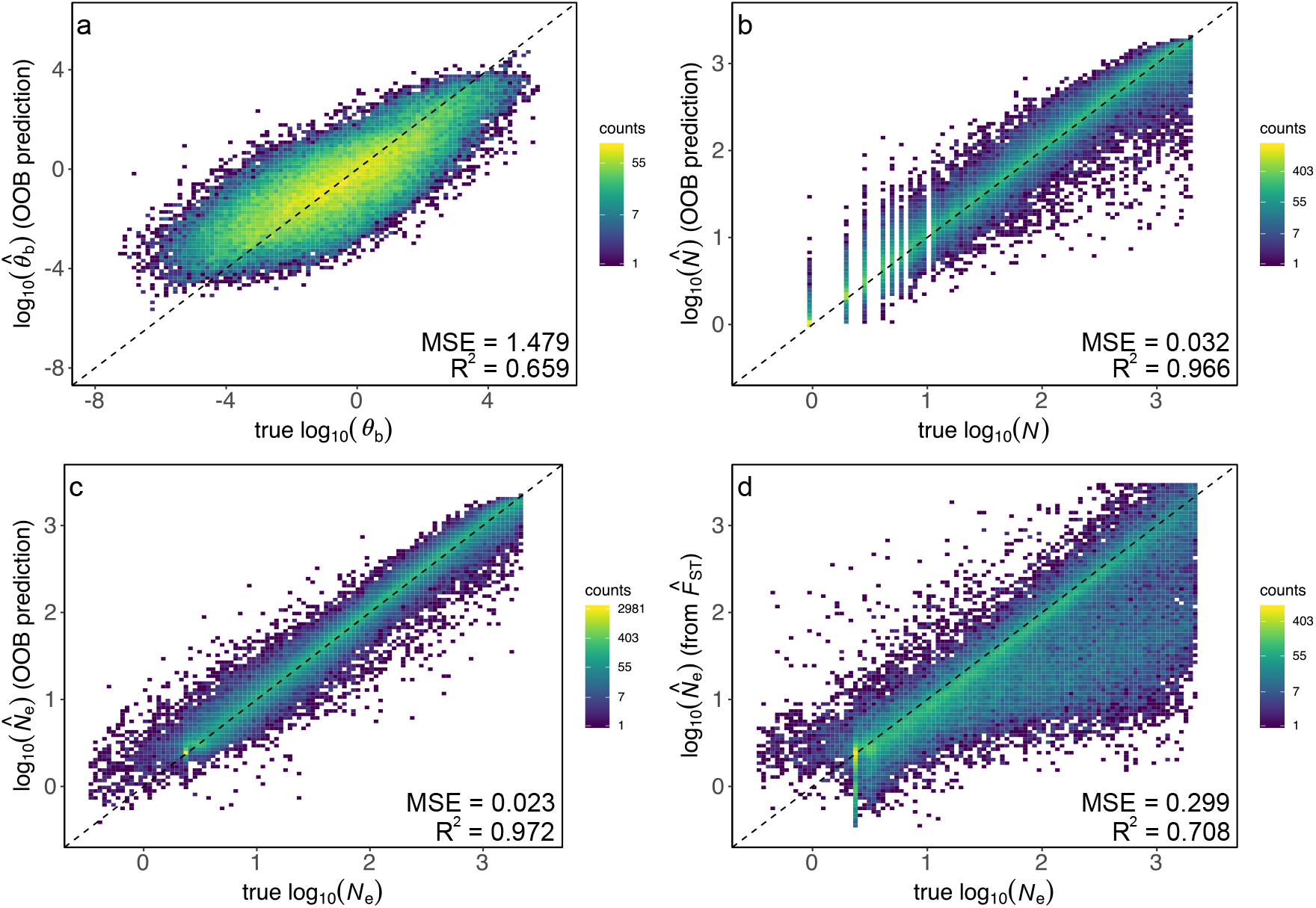
Out-of-bag estimates of ABC-RF trained for the joint inference of demography and selection, and 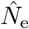 estimates from the temporal *F*_ST_ to compare with the ABC-RF-based 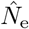 estimates. (a) population scaled mutation rate of beneficial mutations *θ*_b_; (b) population census size *N*; (c) effective population size *N*_e_; and (d) *N*_e_ from temporal *F*_ST_

The automatic selection of informative summary statistics is an important feature of ABC-RF. For each tree of a random forest, summary statistics were selected given its ability to split the data. How many times summary statistics were selected in each RF informs us of their importance for predicting a given parameter. For the prediction of *θ*_b_ values, the RF picked more frequently statistics that reflect the heterogeneity of the genome, such as the 5% quantile of Tajima’s *D* calculated in the second sample, with the kurtosis and skewness of *F*_ST_ and *D*_a_ calculated globally (Figure S6 e). The population size was trained with a combination of within and between sample summary statistics: *F*_ST_ and *D*_a_, with their respective derived statistics frequently selected (Figure S7 c). For *N*_e_, summary statistics that inform about the cumulative divergence between samples as *F*_ST_ and *D*_a_, were frequently selected (Figure S7 d).

### Comparison with *F*_ST_ method to estimate *N*_e_

We compared our ABC-RF *N*_e_ estimates the temporal *F*_ST_ estimates (Frachon et al., 2017). The *F*_ST_-based 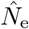 was more affected by the amount of selection in larger populations when selection is more efficient. The *F*_ST_-based 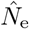 showed higher overall MSE compared to the ABC-based estimates (Figure 2c and d). ABC-RF and the temporal 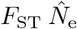 performed well and similarly, regardless of the strength of selection, when the beneficial mutations were less frequent (low *θ*_b_). However, the ABC-based estimator had less local MSE than the temporal *F*_ST_-based estimator. When the frequency of selection is high, the *N*_e_ estimator based on the temporal *F*_ST_ had dramatically higher error (Figure 3).

**Figure 3.**
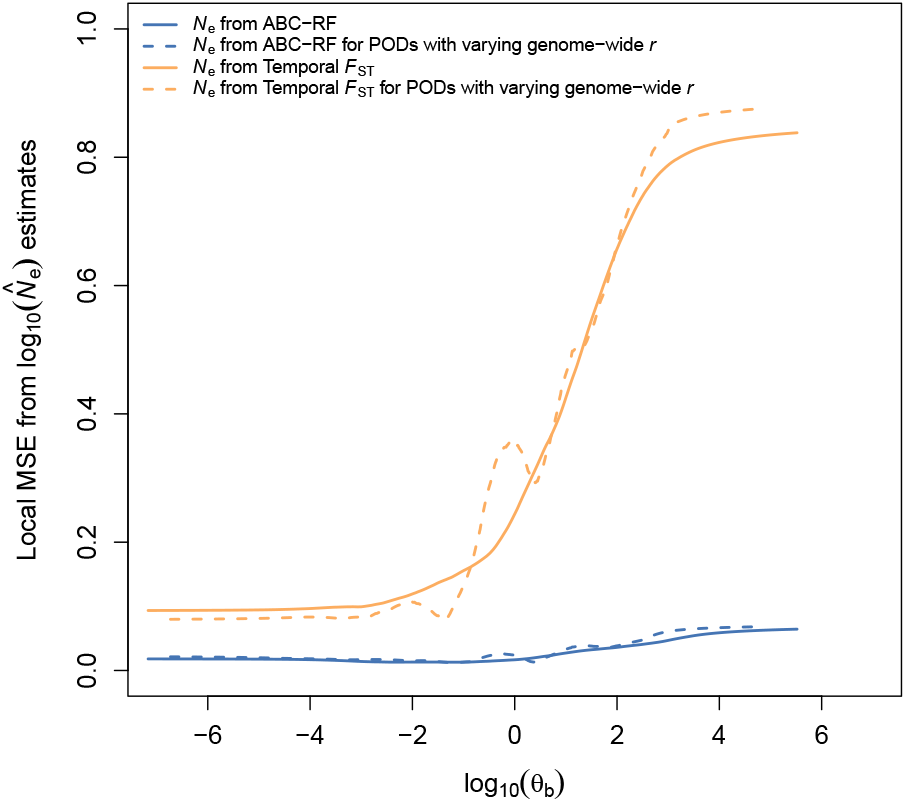
Local MSE of *N*_e_ estimates as a function of *θ*_b_. The lines corresponds to the MSE on *N*_e_ estimates from ABC-RF and from temporal *F*_ST_. Dashed lines correspond to local MSE estimated from pseudo-observed data (POD) with heterogeneous recombination rates along the genome.

### Analysis of temporal genomic data of feral populations of *Apis mellifera*

The projection of each population target data point (in black) into the cloud of the training data points (in grey) in the PCA plots revealed that each population model could capture some dimension of the observed genetic diversity (Figures S8-S14). However, some PCs showed the observed data point outside the simulated data cloud of points, indicating some model inadequacies, possibly because we did not include gene flow or admixture in our simulations. For the analysis of feral *A. mellifera* populations, we first grew independent RF for each parameter in each population. Despite the differences in time intervals between samples, all populations had a similar performance of the ABC-RF estimator for *N*_e_, as they showed similar values of MSE and *R*^2^ (Figure S15). For *N*, trained RF for Humboldt, Stebbins and Placerita performed similarly well, with the lowest MSE and higher *R*^2^ (Figure S16). For *θ*_b_, Riverside had trained RF with the worst performance (Figure S17). Overall, both MSE and *R*^2^ obtained with OOB estimates from simulated data for *A. mellifera* dataset were comparable to these parameters obtained with OOB estimates for the simulated data in the evaluation of the method.

Trained RFs for *N* and *N*_e_ were able to predict these parameters in all populations, as the inference of the mean posterior value and the posterior distribution differentiated from the mean prior value and distribution (Figure 4 b and c). For *N*, posterior distribution were wider than for *N*_e_. Trained RF for *θ*_b_, for all populations had a similar posterior mean, except for the Avalon population that had a peak at a lower value (Figure 4 a). However, the posterior distributions were wider and followed the prior distribution, making it difficult to predict the posterior mean and variance in all populations accurately. It is possible to see together with the posterior mean estimates that the ABC-RF estimates for *θ*_b_ were concentrated in lower values (Table S3) in all populations. *N*_e_ were also lower, and *N*_e_ and *N* were similar. For the results of OOB estimates of other model parameters and latent variables and posterior estimates for these parameters, see section S2 Supplementary Results.

**Figure 4.**
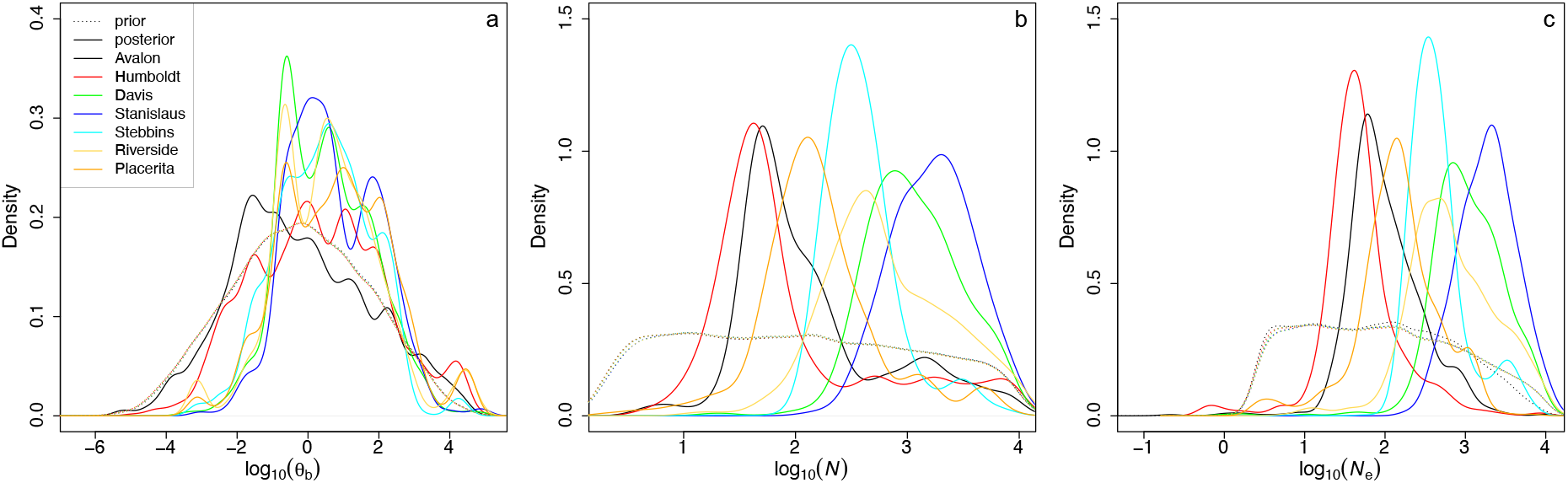
Inference of demography and selection for feral *A. mellifera* populations. (a) the scale mutation rate of selected mutations *θ*_b_, (b) the population census size *N*, (c) the effective population size *N*_e_. Dashed and filled lines correspond to the prior and posterior distributions, respectively. See Table S3, Supplementary Results for mean and 95% credibility intervals.

## Discussion

### Separating demography from drift, and the inference of *θ*_b_

With temporal population genomics data, we can see the evolution in “action” as opposed to single timepoint population genomics data (Feder, PS Pennings, et al., 2021). Consequently, temporal data have more information about the ongoing process, making them interesting for understanding the short-term effects of the interaction between demography and selection (Buffalo and Coop, 2019; Dehasque et al., 2020; Williams and P Pennings, 2020). When samples from more than two time points are available, correlations among allele frequency changes allow to separate the effects of drift and selection (e.g., Buffalo and Coop, 2020; Feder, Kryazhimskiy, et al., 2014). Our results showed that two samples collected at different time points are sufficient for the inference of the genome-wide footprint of adaptive evolution and to separate the demography (population census size *N*) from drift (effective population size *N*_e_).

It is important to stress that *N*_e_, calculated as a latent variable, captures the feedback dynamics between drift and linked selection. Selection, either positive or negative, causes a deviation of *N*_e_ from *N*. The impact of selection on the genome can extend far from the target of selection since individuals that carry beneficial mutations have more chance to reproduce, and their beneficial mutations are more likely to be in the nextgeneration offspring (Walsh and Lynch, 2018). In this complex dynamic, with many loci under selection which creates a dynamic that cannot be easily described analytically, latent variables obtained from simulations can summarize the by-product of drift and selection interactions. With our approach, 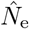 quantifies the drift due to demographic and selection processes, unaffected by the bias of outlier loci.

This genome-wide reduction in *N*_e_ is not captured when loci are assumed to evolve independently (as in Laval et al., 2019; Sheehan and Song, 2016, for example). In contrast, the complexity of linked selection and the genome-wide effect of selection are taken into account using individual-based simulations with the whole genome in an ABC approach.

Estimates of genetic load or other genome-wide parameters about selection are obtained when annotated genomic data is available (Henn et al., 2015), or by conducting experiments on crossing populations (for the genetic load; Plough, 2016). However, we obtained estimates of selection parameters only using polymorphism data. Differently, Buffalo and Coop (2020) measured the genome-wide signature of selection by estimating the covariance of allele frequencies at consecutive time points. The allele frequency covariance matrix allowed the quantification of the genome-wide contribution of selection to the observed allele frequency changes, even when selection involved many loci of small effect. In this work, we estimated the population scale mutation rate of beneficial mutations *θ*_b_, which informs about the diversity of beneficial mutations that existed in the population between the two time points and the potential speed of adaptation at the genome level (Hermisson and PS Pennings, 2017). These estimates reflect the potential number of beneficial mutations between the two time points regardless of their impact as determined by their selection coefficients.

The variable importance plot of each parameter shows us the global importance of each summary statistic in the trained Random Forests. For *N*_e_, *N*, and *θ*_b_ summary statistics calculated from the distribution of locus-specific summary statistics-skewness, kurtosis, mean, variance, 5% and 95% quantiles were more frequently used. Summary statistics derived from the distribution of locus-specific calculated from all segregating loci in the genome inform about the heterogeneity that selection and drift produce genome-wide. For example, a *de novo* a beneficial mutation entered the simulation and was selected; it left a signal of lower diversity around the region it was located. The genome, after selection, contained spots where diversity was high and where it was low, and this heterogeneity was captured by the distribution of locus-specific *H*_E_, more specifically, the lower tail of the distribution where the diversity values of the statistic were lower. The covariance matrix of allele frequencies through time (Buffalo and Coop, 2020) can be used as a summary statistic to capture additional information about the selection and drift when more than two temporal samples are available. Including this matrix as summary statistics for further development of the method would be interesting.

### Comparison with *F*_ST_ method to estimate *N*_e_

We compared the *N*_e_ obtained with ABC-RF framework to the *N*_e_ obtained with *F*_ST_ estimator (Frachon et al., 2017). Overall, the *F*_ST_-based *N*_e_ estimator performed poorly compared to the ABC-RF-based estimator. The lower performance was caused by *N*_e_ values that were underestimated when beneficial mutations were more frequent (higher *θ*_b_). Consequently, the *N*_e_ estimates from the temporal *F*_ST_ were strongly affected by selection. Both estimators performed similarly when the selection was infrequent or rare, but the ABC-RF estimator had lower MSE than the *F*_ST_ one. Positive selection can increase the variance of allele frequency between samples taken at different time points. When selection is infrequent or rare, drift determines most allele frequency changes between samples. Still, when selection is pervasive, selection dominates, which causes dramatic and rapid changes in allele frequency, increasing the variance between samples. *N*_e_ estimator based on the *F*_ST_ depends on the differences in allele frequencies between samples; consequently, it is naturally biased by strong and frequent selection. We can assume that the *N*_e_ estimator from ABC-RF was insensitive to the amount of selection since we trained the ABC-RF with *N*_e_ values from the simulation. In our simulations, *N*_e_ was a latent variable that captured the deviation that selection imposed on the number of individuals able to reproduce (selected for); unaccounted factors did not bias it.

The amount of selection for *θ*_b_ > 1 could be unrealistic in some organisms, but plausible in virus (Feder, Kryazhimskiy, et al., 2014) and many arthropod species, with large *N*_e_, which have larger population sizes (except in eusocial insects that have vertebrate-like population sizes; Romiguier et al., 2014). The selection also acts on weaker and milder beneficial mutations in larger populations. In those organisms, it might be unreasonable to assume mutation-drift equilibrium given the pervasive role of selection. Consequently, attempts to estimate demography parameters as *N*_e_ without properly accounting for the pervasive role of selection could be biased.

### Analysis of temporal genomic data of feral populations of *Apis mellifera*

Overall, the performance of the ABC-RF for selection and demography inference was similar across populations despite the differences in sample size and age. For *θ*_b_, Avalon and Humboldt populations had posterior probability distributions similar to the prior, indicating that the analysis provides no additional information on this parameter. These two populations also present low effective population size estimates, reducing the selection signal. For the rest of the populations, the posterior probability distribution of *θ*_b_ is tilted toward the higher values but without a clear peak differentiating the distribution from the prior. Still, lower *θ*_b_ values could be excluded. It favors the interpretation that selection was acting during the study period without providing a precise parameter estimate. The presence of selection in these analyses comes mainly from the heterogeneity of the polymorphism along the genome. Thus, for a thorough interpretation of the results, it is important to discuss other processes that have not been modeled but could affect this signal. The studied bee populations in California show a mixture of Eastern and Western European ancestry, with some populations presenting African ancestry in the most modern samples Cridland et al. (2018). Different levels of African admixture along the genome could create some heterogeneity and affect the inference. However, in Placerita and Riverside, the populations with higher African ancestry present similar estimates of *θ*_b_ that populations with little or no African admixture. The Humboldt population changed from predominately Western European to Eastern European ancestry, meaning that there was substantial gene flow into the population. These results suggest that admixture does not dramatically affect the inference of selection but also highlights the importance of incorporating admixture in the future development of the approach. Other processes, such as recombination and mutation rate, might be heterogeneous along the genome. Our analysis of simulations with heterogeneous recombination rate suggest that the approach is robust to those. However, more complex models also seem necessary to fully capture the observed genetic diversity (see Figures S7-S13, section S2 Supplementary Results). Including additional factors (admixture, heterogeneity of recombination and mutation rate, and other forms of selection) could be key to obtaining models that fit the data better. Further developments of this approach should take them into account.

Our ABC-RF approach estimated *N*_e_ with the same order of magnitude of other *N*_e_ estimates obtained for hymenopterans (Zayed, 2004). Lower values of *N*_e_ might reflect the presence of admixture, either African admixture or admixture that occurred with domesticated lineages facilitated by changes in beekeeping practices (Cridland et al., 2018). Northern populations, especially from Humboldt County, shared similarities with bees from reared colonies (with higher Eastern European ancestry). Southern populations, as shown by Cridland et al. (2018), showed a higher level of admixture with African lineages. Populations from the southernmost cities (Riverside County, Placerita, and Avalon, Los Angeles County) showed higher genetic diversity than the others but did not show the highest values of *N*_e_. On the other hand, the population of Stanislaus County had the highest value of *N*_e_, possibly because it had lower levels of admixture with domesticated lineages compared to the population from Riverside, Placerita, Avalon, and Los Angeles counties.

We observed that *N*_e_ and *N* had similar estimates. We were aware that our simulation model did not account for key characteristics of eusocial insect reproductive biology: the monopolization of reproduction by the queen and the division of labor. In honey bees, a queen mates with more than one male (a process called polyandry), which leads to a biased breeding sex ratio (Estoup et al., 1994). Assuming that only queens can reproduce in the colony, polyandry increases the variance in the number of parents contributing to the offspring gene pool, which leads to a decrease in the *N*_e_ compared to *N* (Nomura and Takahashi, 2012). In our simulations, we only simulated panmictic random mating. Therefore, the difference between estimates of *N*_e_ and *N* only reflects the selection action. Therefore *N* must be interpreted with caution as it is probably reflecting more the total number of female breeders per generation rather than the size of the population. Individual-based forward simulators such as SLiM allows setting different mating schemes. It is possible to simulate the haplodiploidy, the cast system, diocy, and sex ratio found in honey bees. These simulation modifications could allow us to estimate *N* and other parameters that could better reflect the species’ biology, but that was not the focus of this work.

One possible explanation for the similarities between *N*_e_ and *N* estimates, thus, relies on cast specialization and concentration of reproduction to one of few females in the colony. These came to a cost of reduced *N*_e_, which reduces the efficacy of selection (either positive or negative). Bees are the few insect groups that show very small *N*_e_ potentially linked with the evolution of eusociality (Romiguier et al., 2014). Knowing that lower *N*_e_ reduces the effectiveness of selection, it is plausible to think that lower *N*_e_ is restricting the effects of mutation affecting fitness to stronger beneficial mutations. Since these mutations are less frequent than weak or mild mutations, their effects on *N*_e_ were small, which explains why *N*_e_ and *N* had values in the same range. Low *N*_e_ and low *θ*_b_ pointed to a biological system limited where adaptation is limited by the influx of adaptive mutations (Rousselle et al., 2020).

Our ABC-RF framework also estimated the per-site mutation rate per generation *μ* (Supplementary Results, S18). The mean posterior *μ* for all populations exceeds the mean prior *μ*. The higher estimated values we obtained might be due to the higher true mutation rate but also reflect recent admixture events between these populations. Modeling admixture could help us correctly separate the effects of selection and drift since the introgression of African genes might have biased some estimates of selection parameters.

### Perspectives and Limitations

Our model is relatively simple, as it only considered the impact of new beneficial mutations, neglecting the effect of background selection and standing variation. Background selection can mimic directional selection because it causes a similar pattern of diversity reduction around the target of selection (Stephan, 2010). This has been discussed for a long time; however, much less has been discussed about the patterns of background selection on temporal data and their differences with selective sweeps. Cvijović et al. (2018) showed that neutral alleles linked to less deleterious backgrounds could quickly rise to high frequencies due to purifying selection, which could mimic the signal of a selective sweep in temporal data. However, Schrider (2020) suggests that the signal of selective sweeps will be distinct from the effects of background selection if background selection is not localized to specific regions of the genome. In an attempt to jointly accommodate the effect of demography and selection on the inference of *N*_e_,Johri et al. (2020) modeled the effect of background selection and developed an ABC-based approach that jointly estimated the distribution of fitness effects and *N*_e_. In their simulations, they modeled deleterious mutations and the classical hard sweep with the inclusion of beneficial mutations. They showed an unbiased estimate of *N*_e_ regardless of positive and negative selection presence. Future developments of our approach should include a more realistic genomic architecture were negative and positive mutations can co-occur and explore different concentrations of deleterious mutations.

This will elucidate the importance of background selection in this context, which probably affects our inferences in a way that is difficult to predict with our current results. In addition, further developments should explore scenarios of *de novo* mutations and selection acting on standing variation, which will allow for a more general treatment including soft sweeps. The model can also be expanded to more complex demographic scenarios, including changes in population size and genetic exchange with external sources (migration). Including such admixtures will be key in the future development of this approach since it is also a source of heterogeneity in the genome and, thus, might influence the method’s performance.

## Conclusion

We show that an ABC-RF-based approach can jointly infer adaptive and demographic history from temporal population genomics data. This approach quantifies the genome-wide footprint of selection expressed in the scaled mutation rate of beneficial mutations. The ABC-RF *N*_e_ is robust to varying degrees of strength of selection and frequency of beneficial mutations. Our ABC-RF-based approach can be applied to temporal population genomics datasets to gain insight into natural populations’ adaptive and demographic history.

## Acknowledgements

We are thankful for the Genotoul bioinformatics platform Toulouse Occitanie (Bioinfo Genotoul, https://doi.org/10.15454/1.5572369328961167E12), for the High-Performance Computing Center University of Montpellier (MESO@LR-Platform) and for the Ohio Supercomputer Center (Ohio Supercomputer Center, 1987) for providing computing resources. The authors would like to thank Julie Cridland for sharing the processed data and Andrew P. Michel for providing suggestions to improve an early version of the manuscript. Preprint version 6 of this article has been peer-reviewed and recommended by Peer Community In Evolutionary Biology (https://doi.org/10.24072/pci.evolbiol.100158).

## Data, script, code, and supplementary information availability

Data are available online: https://doi.org/10.5281/zenodo.4599735 Script and codes are available online: https://doi.org/10.5281/zenodo.4599735 Supplementary information is available online: https://doi.org/10.1101/2021.03.12.435133

