## Supplementary Material for "Joint inference of adaptive and demographic history from temporal population genomic data"

November 8, 2022

### S1 Supplementary Methods

#### S1.1 List of summary statistics

Three groups of summary statistics were calculated for the reference table (e.g. training data): 1) summary statistics calculated SNP-by-SNPs (i.e., locus-specific summary statistics - LSS); 2) summary statistics calculated for nucleotide sequence (i.e., window summary statistics - WSS) - in this case, a window containing base positions was defined around each SNP in the dataset, and 3) summary statistics calculated globally by averaging the locus-specific or window-specific summary statistics (global summary statistics - GSS). Summary statistics about within-population diversity as $H_{\mathrm{E}}$ were calculated for each time point individually and marginally by pooling all individuals sampled together. Three window sizes were used for the summary statistics calculated in a window: 500bp, 5Kbp, and 10Kbp. We used the allele counts instead of frequency for the site-frequency spectrum, and all bins were included in the reference table.

1. LSSs - Locus-specific summary statistics:
   (a) $D_{\mathrm{J}}$ - Jost's $D$ (Jost, 2008, 2009);
   (b) $F_{\mathrm{ST}}$ - Weir and Cockerham's $F_{\mathrm{ST}}$ (Weir and Cockerham, 1984);
   (c) $H_{\mathrm{E}}$ - Expected Heterozygosity;
2. WSSs - Window-specific summary statistics (window sizes - 500bp, 5Kbp and 10Kbp):
   (a) $D_{\mathrm{j}}$ - Jost's $D$;
   (b) $F_{\mathrm{ST}}$ - Weir and Cockerham's $F_{\mathrm{ST}}$;
   (c) $D_{\mathrm{a}}$ - Net distance between populations;
   (d) $H_{\mathrm{E}}$ - Expected Heterozygosity;
   (e) $S$ - Number of polymorphic sites;
   (f) $\pi$ - Nucleotide diversity (Nei and Li, 1979);
   (g) $\theta_{\mathrm{W}}$ - Watterson's 4Nu estimator (Watterson, 1975);
   (h) $D$ - Tajima's D (Tajima, 1989);
3. Global summary statistics:
   (a) All summary statistics enumerated above (including the within population calculations);
   (b) SFS bins - binned Joint Site-frequency spectrum $SFS$ (Ewing and Jensen, 2016);
   (c) mean, variance, kurtosis, skewness, and the 5% and 95% quantiles of all intra-locus summary statistics presented above (LSSs and WSSs);

### S2 Supplementary Results

#### S2.1 ABC-RF

Figure S1 shows the prior distribution of the parameters and latent variables estimated with the ABC-RF. These distributions correspond to the values that were used to train ABC-Random forests.

Figure S2 shows the distribution of Random Forest OOB estimates on the observed values (the ones used to train the RF) for model parameters and other latent variables informative about selection. The ABC-RF regressions performed poorly on the inference of two parameters of the selection dynamics $P_{\mathrm{R}}P_{\mathrm{B}}$ and $\gamma$. On the other hand, the ABC-RF performed better on the latent variables $P$ and $\bar{s}$ (which can be viewed as the realization of the two parameters mentioned above). Each value of these latent variables was obtained from the simulation by 1) calculating the relative proportion of strongly beneficial mutations (those with $s > 1/N_{\mathrm{e}}$) over all possible segregating mutations, 2) averaging their $s$ values.

The estimation of substitution load $L$ also had a somehow good performance. It performed well for higher values but poorly for lower ones, as shown in the distribution of the estimated OOB values against the true values (Figure S2 e). Only strong adaptive dynamics (many loci under selection or strong selection) seem to leave informative polymorphism patterns about substitution load. The proportion of strongly beneficial mutations that were segregating in the population, $P$, can also be used as a complementary estimator to characterize the adaptive dynamics (it is complementary to $\theta_{\mathrm{b}}$ presented in the main text).

Figure S3 also presents the distribution OOB estimates with the true observed values for model parameters and the latent variables informative about drift. ABC-RF could not satisfactorily recover the recombination rate $r$ as expected for these model parameters, given the calculated summary statistics and the information contained in the temporal allele frequency dynamics. On the other hand, the ABC-RF estimated the mutation rate per nucleotide position per generation $\mu$ satisfactorily. The last latent variable also performed relatively well and similarly to the $L$. It could be used with other estimates to characterize the selection/drift dynamics when selection is more frequent.

To test the robustness of our ABC-RF-based framework, we applied the trained RFs on PODs with across-genome varying recombination rates. Figure S4 depicts how recombination rates were set on simulations. Simulations for ABC-RF training had fixed genome-wide recombination rates per generation $r$; Additional 1,000 simulations for testing the framework's robustness had varying recombination rates. The scatterplots showed a similar dispersion pattern as the OOB plots when the true values were plotted against the estimated values (Figure S5). Quantitatively, these PODs presented similar behavior regarding the MSE and $R^2$ estimates (Figure S5, and Table S2).

Table S2 contains MSE and bias for $P$, $\bar{s}$, and $L$ from neutral simulations. For training simulations (for model proof of concept and analysis of the feral bee populations), we first removed neutral simulations for these parameters (where values were zero) before RF training. Then each trained RF was used to make predictions on these neutral simulations. The MSE and bias correspond to the difference between the true values (zeros) and the predicted values in the original scale. For simulations used to test the robustness of our framework when there are varying rates of recombination, we calculated MSE and $R^2$ for non-neutral PODs (Figure S5) and MSE and bias for neutral PODs.

Figure S6 and Figure S7 show the variable importance plots (VIP) of each ABC-RF that were grown to infer the parameters of our ABC-RF framework. In a nutshell, the VIP shows how many times the RF selected summary statistics during the training phase. Consequently, the VIP shows the relative importance of each summary statistic for training that parameter (more details in the main text). For the latent variables that account for the selection and drift dynamics, the top summary statistics were those calculated from the genome-wide distribution of locus-specific summary statistics that reflect their genome-wide distribution - variance, mean, skewness, kurtosis, and quantiles. RF frequently used summary statistics that inform about the distribution of locus-specific heterozygosity, calculated globally and calculated for the second sample. And also, summary statistics informative about the distribution of genome-wide intra-locus $F_{\mathrm{ST}}$, for the latent variables $P$ and $\bar{s}$. For $L$ summary statistics informative about the genetic divergence between samples: $F_{\mathrm{ST}}$, Da and Dj were more frequently selected.

#### S2.2 Analysis of temporal genomic data of feral populations of *Apis mellifera*

Before we trained the RF to analyze feral bee populations, we checked if our model could capture most of the features of the observed genomic data (target dataset). We run a principal component analysis (PCA) and projected pairs of principal components (PC) on a PCA plot for each population reference table containing only the summary statistics from the simulated data. We also ran a PCA on the observed (or target) dataset summary statistics and projected the same PCs on the PCA plot computed with simulated data. The model mostly capture features of the data, however some PCs showed the observed data point (in black) far from the simulated data cloud of points (in grey), indicating some model inadequacies possibly due to the fact that we did not include gene flow and/or admixture in our simulations (Figure S8, Figure S9, Figure S10, Figure S11, Figure S12, Figure S13, Figure S14).

Figure S17, Figure S16, and Figure S15 show the distribution of the true and the OOB estimated values for the parameters used in this work for the joint inference of demography and selection for temporal pairs of populations of feral *A. mellifera.* The OOB MSE and $R^2$ estimates showed that even with fewer simulations for model training, the RF grew for the *A. mellifera* data had comparable performance to the RF grew for the simulated data.

The performance of the model parameters and additional latent variables for all temporal pairs of this dataset were similar to the simulations used for the framework evaluation. Figure S18, Figure S19, Figure S20, Figure S21, and Figure S22 show the parameters and latent variables informative about the adaptive history. Figure S23, Figure S24, and Figure S25 show the model parameters and an additional latent variable complementary to the mutation load for the inference of the selection impact on demography. For comparison purposes, we also included the scatterplots of true $N_{\mathrm{e}}$ against the $F_{\mathrm{ST}}$-NE computed on the global $F_{\mathrm{ST}}$ of simulated data for all feral bee populations (Figure S26).

Table S3 contains the posterior estimates of each parameter for each *A. mellifera* population. The last row shows the $F_{\mathrm{ST}}$-NE estimates for comparison. Figure S27 and Figure S28 show the prior, and posterior distributions of all remaining model parameters and latent variables calculated for the temporal samples analyzed with the ABC-RF framework.

**Table S1: Notation.**

| Symbol | Meaning |
|---|---|
| $r$ | recombination rate per base pair per generation |
| $G$ | genome size |
| $h$ | dominance coefficient |
| $L$ | mean substitution load |
| $N$ | population size during the inference period |
| $N_0$ | population size during the burn-in period |
| $N_\mathrm{e}$ | effective population size during the inference period |
| $P_\mathrm{R}$ | proportion of non-neutral regions |
| $P_\mathrm{B}$ | proportion of beneficial mutations per non-neutral region |
| $P$ | proportion of polymorphisms under strong selection, i.e. $s > 1/N_\mathrm{e}$ |
| $s$ | selection coefficient |
| $\bar{s}$ | average selection coefficients of mutations under strong selection |
| $t_1$ and $t_2$ | generations were samples of individuals were taken |
| $W_{\mathrm{max}i}$ | fitness of individual with highest fitness in the population in generation $i$ |
| $\bar{W}_i$ | mean fitness of the population in generation $i$ |
| $\gamma$ | mean of the gamma distribution where the selection coefficients of each beneficial mutation was sampled |
| $\mu$ | mutation rate per generation |
| $\mu_\mathrm{b}$ | mutation rate of beneficial mutations per generation |
| $\tau$ | length of time-interval between the first and the second sample |
| $\theta_\mathrm{b}$ | scaled mutation rate of beneficial mutations |

**Table S2: Neutral simulations' MSE and Bias.**

| Parameter | Dataset | MSE | Bias |
|---|---|---|---|
| | Training | 0.001 | 0.014 |
| | PODs recombination | 0.001 | 0.013 |
| | Avalon | 0 | 0.007 |
| | Humboldt | 0 | 0.009 |
| $P$ | Davis | 0 | 0.009 |
| | Stanislaus | 0 | 0.010 |
| | Stebbins | 0 | 0.013 |
| | Riverside | 0 | 0.013 |
| | Placerita | 0 | 0.013 |
| | Training | 0.262 | 0.401 |
| | PODs recombination | 0.241 | 0.386 |
| | Avalon | 0.238 | 0.410 |
| | Humboldt | 0.318 | 0.471 |
| $\bar{s}$ | Davis | 0.305 | 0.467 |
| | Stanislaus | 0.315 | 0.479 |
| | Stebbins | 0.373 | 0.533 |
| | Riverside | 0.368 | 0.540 |
| | Placerita | 0.379 | 0.542 |
| | Training | 0.014 | 0.089 |
| | PODs recombination | 0.013 | 0.088 |
| | Avalon | 0.011 | 0.031 |
| | Humboldt | 0.006 | 0.024 |
| $L$ | Davis | 0.010 | 0.034 |
| | Stanislaus | 0.007 | 0.032 |
| | Stebbins | 0.008 | 0.033 |
| | Riverside | 0.011 | 0.046 |
| | Placerita | 0.008 | 0.034 |

**Table S4: Summary statistics for *A. mellifera* feral populations.**

| | Avalon | Humboldt | Davis | Stanislaus | Stebbins | Riverside | Placerita |
|---|---|---|---|---|---|---|---|
| GSS $H_{\mathrm{E}}$ | 0.37 | 0.29 | 0.28 | 0.28 | 0.25 | 0.22 | 0.21 |
| GSS $D_{\mathrm{J}}$ | 0.17 | 0.12 | 0 | 0 | 0 | 0 | 0.02 |
| GSS $F_{\mathrm{ST}}$ | 0.28 | 0.26 | 0.02 | 0.01 | 0.01 | 0.01 | 0.05 |
| GSS $S$ | 43187 | 61431 | 51268 | 61264 | 69961 | 113897 | 98438 |
| GSS $\theta_{\mathrm{W}}$ | 13916.88 | 16450.51 | 16121.33 | 18462.86 | 19719.88 | 32104.1 | 27746.68 |
| GSS $\pi$ | 16106.09 | 17784.77 | 14465.06 | 16938.75 | 17257.82 | 24881.59 | 20222.57 |
| GSS $D$ | 0.73 | 0.33 | -0.47 | -0.36 | -0.52 | -0.94 | -1.14 |
| GSS $D_{\mathrm{a}}$ | 0.14 | 0.08 | 0 | 0 | 0 | 0 | 0.01 |
| GSS $t_1$ $H_{\mathrm{E}}$ | 0.15 | 0.26 | 0.29 | 0.29 | 0.26 | 0.2 | 0.15 |
| GSS $t_2$ $H_{\mathrm{E}}$ | 0.36 | 0.23 | 0.28 | 0.27 | 0.23 | 0.22 | 0.23 |
| GSS $t_1$ $S$ | 8280 | 46969 | 27675 | 32530 | 54885 | 43351 | 40960 |
| GSS $t_2$ $S$ | 36347 | 45561 | 43311 | 54571 | 48762 | 103077 | 83254 |
| GSS $t_1$ $\theta$W | 5520 | 15553.28 | 15095.45 | 17743.64 | 19401.07 | 23646 | 15797.25 |
| GSS $t_2$ $\theta$W | 12848.15 | 15087.04 | 15309.82 | 18070.6 | 17236.67 | 31063.86 | 27568.67 |
| GSS $t_1$ $\pi$ | 5592 | 16163.6 | 15055.33 | 17661 | 18281.3 | 23115.33 | 14802.06 |
| GSS $t_2$ $\pi$ | 15535.81 | 14006.38 | 14116 | 16630.37 | 15978.22 | 25182.07 | 22708.71 |
| GSS $t_1$ $D$ | 0 | 0.19 | -0.03 | -0.05 | -0.29 | -0.24 | -0.35 |
| GSS $t_2$ $D$ | 1.05 | -0.34 | -0.39 | -0.38 | -0.37 | -0.83 | -0.83 |
| SFS $\mathrm{bin}_1$ | 0 | 16110 | 0 | 0 | 25318 | 48723 | 39536 |
| SFS $\mathrm{bin}_2$ | 9364 | 7870 | 20589 | 22294 | 12257 | 23856 | 24263 |
| SFS $\mathrm{bin}_3$ | 3403 | 5514 | 11095 | 11758 | 8423 | 12827 | 11460 |
| SFS $\mathrm{bin}_4$ | 4781 | 4394 | 182 | 8575 | 6313 | 8346 | 6628 |
| SFS $\mathrm{bin}_5$ | 5623 | 6770 | 7313 | 6212 | 4818 | 5783 | 4684 |
| SFS $\mathrm{bin}_6$ | 4580 | 2931 | 4588 | 6 | 3852 | 4285 | 3159 |
| SFS $\mathrm{bin}_7$ | 3895 | 3319 | 72 | 4694 | 3062 | 3419 | 2852 |
| SFS $\mathrm{bin}_8$ | 4016 | 3801 | 3380 | 3428 | 2539 | 2997 | 2331 |
| SFS $\mathrm{bin}_9$ | 4572 | 4094 | 2752 | 2901 | 2280 | 2509 | 2159 |
| SFS $\mathrm{bin}_1 0$ | 2953 | 6628 | 1297 | 1396 | 1099 | 1152 | 1366 |
| LSS mean($H_{\mathrm{E}}$) | 0.37 | 0.29 | 0.28 | 0.28 | 0.25 | 0.22 | 0.21 |
| LSS mean($D_{\mathrm{J}}$) | 0.17 | 0.12 | 0 | 0 | 0 | 0 | 0.02 |
| LSS mean($F_{\mathrm{ST}}$) | 0.15 | 0.14 | -0.01 | -0.02 | 0.01 | -0.02 | 0.02 |

*continued on next page*

*continued from previous page*

| | Avalon | Humboldt | Davis | Stanislaus | Stebbins | Riverside | Placerita |
|---|---|---|---|---|---|---|---|
| LSS $t_1$ mean($H_E$) | 0.15 | 0.26 | 0.29 | 0.29 | 0.26 | 0.2 | 0.15 |
| LSS $t_2$ mean($H_E$) | 0.36 | 0.23 | 0.28 | 0.27 | 0.23 | 0.22 | 0.23 |
| WSS mean($H_E$) | 0.37 | 0.29 | 0.28 | 0.28 | 0.25 | 0.22 | 0.21 |
| WSS mean($D_J$) | 0.17 | 0.12 | 0 | 0 | 0 | 0 | 0.02 |
| WSS mean($F_{ST}$) | 0.23 | 0.22 | 0.01 | 0 | 0.01 | 0 | 0.05 |
| WSS mean($S$) | 7.07 | 9.3 | 8.02 | 9.28 | 10.39 | 16.15 | 14.17 |
| WSS mean($\theta_W$) | 2.3 | 2.49 | 2.52 | 2.8 | 2.93 | 4.55 | 4 |
| WSS mean($\pi$) | 2.64 | 2.7 | 2.28 | 2.57 | 2.57 | 3.53 | 2.91 |
| WSS mean($D$) | 0.53 | 0.26 | -0.37 | -0.29 | -0.42 | -0.82 | -0.97 |
| WSS mean($D_a$) | 0.14 | 0.08 | 0 | 0 | 0 | 0 | 0.01 |
| WSS $t_1$ mean($H_E$) | 0.15 | 0.26 | 0.29 | 0.29 | 0.26 | 0.2 | 0.15 |
| WSS $t_2$ mean($H_E$) | 0.36 | 0.23 | 0.28 | 0.27 | 0.23 | 0.22 | 0.23 |
| WSS $t_1$ mean($S$) | 1.3 | 7.13 | 4.35 | 4.92 | 8.11 | 6.14 | 5.81 |
| WSS $t_2$ mean($S$) | 6.02 | 6.93 | 6.81 | 8.29 | 7.3 | 14.61 | 12.07 |
| WSS $t_1$ mean($\theta_W$) | 0.86 | 2.36 | 2.38 | 2.69 | 2.87 | 3.35 | 2.25 |
| WSS $t_2$ mean($\theta_W$) | 2.13 | 2.29 | 2.41 | 2.74 | 2.58 | 4.4 | 4 |
| WSS $t_1$ mean($\pi$) | 0.88 | 2.45 | 2.37 | 2.67 | 2.71 | 3.28 | 2.1 |
| WSS $t_2$ mean($\pi$) | 2.57 | 2.13 | 2.22 | 2.52 | 2.38 | 3.57 | 3.28 |
| WSS $t_1$ mean($D$) | 0.78 | 0.14 | -0.06 | -0.05 | -0.24 | -0.21 | -0.32 |
| WSS $t_2$ mean($D$) | 0.83 | -0.28 | -0.32 | -0.31 | -0.31 | -0.72 | -0.72 |
| LSS variance($H_E$) | 0.02 | 0.03 | 0.02 | 0.02 | 0.02 | 0.02 | 0.02 |
| LSS variance($D_J$) | 0.08 | 0.04 | 0 | 0.02 | 0 | 0.01 | 0.01 |
| LSS variance($F_{ST}$) | 0.12 | 0.05 | 0.04 | 0.04 | 0.01 | 0.03 | 0.01 |
| LSS $t_1$ variance($H_E$) | 0.09 | 0.04 | 0.08 | 0.08 | 0.03 | 0.07 | 0.04 |
| LSS $t_2$ variance($H_E$) | 0.04 | 0.03 | 0.03 | 0.03 | 0.03 | 0.02 | 0.02 |
| WSS variance($H_E$) | 0.01 | 0.01 | 0 | 0 | 0 | 0 | 0 |
| WSS variance($D_J$) | 0.03 | 0.01 | 0 | 0 | 0 | 0 | 0 |
| WSS variance($F_{ST}$) | 0.05 | 0.02 | 0.02 | 0.02 | 0.01 | 0.01 | 0.01 |
| WSS variance($S$) | 18.52 | 27.66 | 21 | 27.23 | 33.73 | 71.26 | 63.45 |
| WSS variance($\theta_W$) | 1.98 | 1.98 | 2.08 | 2.47 | 2.68 | 5.66 | 5.07 |
| WSS variance($\pi$) | 3.06 | 3.01 | 2.01 | 2.5 | 2.43 | 4.18 | 3.07 |
| WSS variance($D$) | 0.75 | 0.69 | 0.56 | 0.55 | 0.5 | 0.32 | 0.33 |
| WSS variance($D_a$) | 0.02 | 0.01 | 0 | 0 | 0 | 0 | 0 |
| WSS $t_1$ variance($H_E$) | 0.03 | 0.01 | 0.02 | 0.02 | 0.01 | 0.01 | 0.01 |
| WSS $t_2$ variance($H_E$) | 0.01 | 0.01 | 0.01 | 0.01 | 0.01 | 0 | 0 |
| WSS $t_1$ variance($S$) | 1.74 | 20.22 | 10.58 | 12.44 | 22.62 | 18.31 | 15.17 |
| WSS $t_2$ variance($S$) | 15.87 | 20.1 | 17.83 | 23.65 | 21.43 | 60.91 | 51.58 |
| WSS $t_1$ variance($\theta_W$) | 0.77 | 2.22 | 3.19 | 3.7 | 2.82 | 5.45 | 2.29 |
| WSS $t_2$ variance($\theta_W$) | 1.98 | 2.21 | 2.23 | 2.59 | 2.68 | 5.53 | 5.66 |
| WSS $t_1$ variance($\pi$) | 0.85 | 2.61 | 3.21 | 3.7 | 2.92 | 5.26 | 2.17 |
| WSS $t_2$ variance($\pi$) | 3.23 | 2.21 | 2.09 | 2.48 | 2.38 | 4.31 | 4.04 |
| WSS $t_1$ variance($D$) | 1.95 | 0.65 | 0.61 | 0.58 | 0.52 | 0.38 | 0.55 |
| WSS $t_2$ variance($D$) | 0.87 | 0.65 | 0.61 | 0.59 | 0.6 | 0.33 | 0.37 |
| LSS kurtosis($H_E$) | 1.74 | 1.42 | 1.83 | 1.74 | 1.89 | 2.54 | 2.9 |
| LSS kurtosis($D_J$) | 5.1 | 4.6 | 0 | 12.05 | 18.94 | 18.45 | 25.7 |
| LSS kurtosis($F_{ST}$) | 3.04 | 3.47 | 4.88 | 5.17 | 7.23 | 5.61 | 8.51 |
| LSS $t_1$ kurtosis($H_E$) | 4.44 | 1.62 | 1.32 | 1.18 | 1.9 | 1.47 | 2.39 |
| LSS $t_1$ kurtosis($H_E$) | 2.19 | 1.86 | 2.1 | 2.04 | 1.82 | 2.42 | 2.39 |
| WSS kurtosis($H_E$) | 2.98 | 3.35 | 3.42 | 3.43 | 3.83 | 5.5 | 5.32 |
| WSS kurtosis($D_J$) | 6.33 | 6.02 | 0 | 16.68 | 15.31 | 13.31 | 37.84 |
| WSS kurtosis($F_{ST}$) | 3.4 | 2.84 | 6.09 | 5.97 | 6.58 | 5.67 | 6.01 |
| WSS kurtosis($\pi$) | 18.42 | 14.94 | 7.94 | 13.06 | 11.08 | 13.93 | 14.56 |
| WSS kurtosis($D$) | 2.68 | 2.69 | 2.85 | 2.8 | 3 | 3.71 | 3.84 |
| WSS kurtosis($D_a$) | 9.82 | 7.77 | 41.89 | 37.07 | 19.05 | 15.6 | 56.6 |
| WSS $t_1$ kurtosis($H_E$) | 6.45 | 3.34 | 2.68 | 2.8 | 3.7 | 3.94 | 4.49 |
| WSS $t_2$ kurtosis($H_E$) | 3.57 | 3.41 | 3.66 | 3.67 | 3.67 | 5.31 | 5.16 |
| WSS $t_1$ kurtosis($S$) | 5.61 | 12.23 | 8.21 | 8.58 | 9.46 | 13.76 | 10.17 |
| WSS $t_2$ kurtosis($S$) | 13.32 | 13.22 | 10.47 | 9.74 | 12.76 | 8.21 | 17.75 |
| WSS $t_1$ kurtosis($\theta_W$) | 6.46 | 12.23 | 8.66 | 8.58 | 9.46 | 13.76 | 10.35 |
| WSS $t_2$ kurtosis($\theta_W$) | 13.32 | 13.25 | 10.46 | 9.74 | 12.76 | 8.21 | 17.73 |
| WSS $t_1$ kurtosis($\pi$) | 6.52 | 10.15 | 8.33 | 8.19 | 11.37 | 13.94 | 9.96 |
| WSS $t_2$ kurtosis($\pi$) | 18.8 | 12.82 | 8.08 | 9.59 | 8.21 | 12.68 | 15.89 |
| WSS $t_1$ kurtosis($D$) | 5.29 | 2.56 | 4.82 | 3.19 | 2.77 | 4.51 | 2.93 |
| WSS $t_2$ kurtosis($D$) | 2.92 | 2.62 | 2.64 | 2.74 | 2.67 | 3.58 | 3.52 |
| LSS skewness($H_E$) | -0.39 | 0.08 | 0.47 | 0.39 | 0.54 | 0.89 | 1.08 |
| LSS skewness($D_J$) | 1.22 | 1.58 | 0 | 2.09 | 2.95 | 2.59 | 3.8 |
| LSS skewness($F_{ST}$) | 0.78 | 1.14 | 1.39 | 1.58 | 1.65 | 1.81 | 1.88 |
| LSS $t1$ skewness($H_E$) | 1.68 | -0.03 | 0.02 | 0 | 0.01 | 0.58 | 0.88 |
| LSS $t2$ skewness($H_E$) | -0.77 | 0.25 | -0.03 | 0.1 | 0.2 | 0.58 | 0.39 |
| WSS skewness($H_E$) | -0.42 | 0.08 | 0.36 | 0.32 | 0.4 | 0.72 | 0.77 |
| WSS skewness($D_J$) | 1.25 | 1.36 | 0 | 2.08 | 2.18 | 1.51 | 3.19 |
| WSS skewness($F_{ST}$) | 0.51 | 0.44 | 1.37 | 1.4 | 1.47 | 1.16 | 1.21 |
| WSS skewness($\pi$) | 2.48 | 2.28 | 1.57 | 1.98 | 1.83 | 2.11 | 2.18 |
| WSS skewness($D$) | -0.4 | 0.01 | 0.32 | 0.26 | 0.36 | 0.57 | 0.69 |
| WSS skewness($D_a$) | 1.99 | 1.69 | 3.5 | 3.43 | 2.62 | 1.93 | 3.89 |
| WSS $t_1$ skewness($H_E$) | 1.52 | 0 | -0.07 | -0.04 | 0.01 | 0.37 | 0.61 |
| WSS $t_2$ skewness($H_E$) | -0.64 | 0.12 | -0.15 | 0.06 | 0.07 | 0.5 | 0.21 |
| WSS $t_1$ skewness($S$) | 1.34 | 1.96 | 1.58 | 1.64 | 1.63 | 2.06 | 1.79 |
| WSS $t_2$ skewness($S$) | 2.05 | 2.06 | 1.74 | 1.64 | 1.91 | 1.5 | 2.42 |
| WSS $t_1$ skewness($\theta_W$) | 1.41 | 1.96 | 1.65 | 1.64 | 1.63 | 2.06 | 1.8 |
| WSS $t_2$ skewness($\theta_W$) | 2.05 | 2.07 | 1.74 | 1.64 | 1.91 | 1.5 | 2.42 |
| WSS $t_1$ skewness($\pi$) | 1.48 | 1.8 | 1.62 | 1.61 | 1.91 | 2.08 | 1.79 |
| WSS $t_2$ skewness($\pi$) | 2.49 | 2.12 | 1.54 | 1.69 | 1.52 | 2.01 | 2.3 |
| WSS $t_1$ skewness($D$) | 1.2 | -0.04 | 1.24 | 1 | 0.26 | 1.29 | 0.36 |
| WSS $t_2$ skewness($D$) | -0.68 | 0.28 | 0.3 | 0.3 | 0.32 | 0.53 | 0.61 |
| LSS quantile(0.05,$H_E$) | 0.14 | 0.08 | 0.14 | 0.13 | 0.1 | 0.1 | 0.09 |
| LSS quantile(0.05, $D_J$) | -0.13 | -0.04 | -0.16 | -0.15 | -0.09 | -0.13 | -0.06 |
| LSS quantile(0.05, $F_{ST}$) | -0.32 | -0.09 | -0.2 | -0.2 | -0.11 | -0.17 | -0.11 |
| LSS $t_1$ quantile(0.05,$H_E$) | 0 | 0 | 0 | 0 | 0 | 0 | 0 |
| LSS $t_2$ quantile(0.05,$H_E$) | 0 | 0 | 0 | 0 | 0 | 0 | 0 |
| WSS quantile(0.05,$H_E$) | 0.22 | 0.17 | 0.17 | 0.17 | 0.15 | 0.15 | 0.13 |
| WSS quantile(0.05, $D_J$) | -0.03 | 0 | -0.09 | -0.08 | -0.04 | -0.04 | -0.02 |
| WSS quantile(0.05, $F_{ST}$) | -0.1 | -0.01 | -0.15 | -0.15 | -0.07 | -0.11 | -0.05 |
| WSS quantile(0.05, $S$) | 2 | 3 | 2 | 3 | 3 | 5 | 4 |
| WSS quantile(0.05, $\theta_W$) | 0.64 | 0.8 | 0.63 | 0.9 | 0.85 | 1.41 | 1.13 |
| WSS quantile(0.05, $\pi$) | 0.55 | 0.6 | 0.53 | 0.63 | 0.59 | 0.96 | 0.74 |
| WSS quantile(0.05, $D$) | -1.1 | -1.15 | -1.51 | -1.48 | -1.51 | -1.66 | -1.8 |

*continued on next page*

*continued from previous page*

| | Avalon | Humboldt | Davis | Stanislaus | Stebbins | Riverside | Placerita |
|---|---|---|---|---|---|---|---|
| WSS quantile(0.05, $D_{\mathrm{a}}$) | 0 | 0 | -0.05 | -0.05 | -0.02 | -0.03 | -0.01 |
| WSS $t_1$ quantile(0.05,$H_{\mathrm{E}}$) | 0 | 0.12 | 0 | 0.06 | 0.13 | 0.06 | 0.04 |
| WSS $t_2$ quantile(0.05,$H_{\mathrm{E}}$) | 0.18 | 0.08 | 0.13 | 0.15 | 0.09 | 0.14 | 0.15 |
| WSS $t_1$ quantile(0.05, $S$) | 0 | 2 | 0 | 1 | 2 | 1 | 1 |
| WSS $t_2$ quantile(0.05, $S$) | 1 | 1 | 1 | 2 | 1 | 4 | 3 |
| WSS $t_1$ quantile(0.05, $\theta_{\mathrm{W}}$) | 0 | 0.66 | 0 | 0.55 | 0.71 | 0.55 | 0.39 |
| WSS $t_2$ quantile(0.05, $\theta_{\mathrm{W}}$) | 0.35 | 0.33 | 0.35 | 0.66 | 0.35 | 1.21 | 0.99 |
| WSS $t_1$ quantile(0.05, $\pi$) | 0 | 0.48 | 0 | 0.5 | 0.57 | 0.5 | 0.25 |
| WSS $t_2$ quantile(0.05, $\pi$) | 0.47 | 0.33 | 0.4 | 0.55 | 0.4 | 0.95 | 0.8 |
| WSS $t_1$ quantile(0.05, $D$) | -0.71 | -1.18 | -0.81 | -0.81 | -1.4 | -0.82 | -1.45 |
| WSS $t_2$ quantile(0.05, $D$) | -1.06 | -1.53 | -1.56 | -1.53 | -1.49 | -1.59 | -1.63 |
| LSS quantile(0.95,$H_{\mathrm{E}}$) | 0.55 | 0.52 | 0.53 | 0.53 | 0.51 | 0.51 | 0.51 |
| LSS quantile(0.95, $D_{\mathrm{J}}$) | 0.89 | 0.59 | 0.28 | 0.18 | 0.09 | 0.13 | 0.15 |
| LSS quantile(0.95, $F_{\mathrm{ST}}$) | 0.84 | 0.59 | 0.3 | 0.32 | 0.22 | 0.37 | 0.25 |
| LSS $t_1$ quantile(0.95,$H_{\mathrm{E}}$) | 1 | 0.53 | 0.67 | 0.67 | 0.53 | 0.67 | 0.54 |
| LSS $t_2$ quantile(0.95,$H_{\mathrm{E}}$) | 0.56 | 0.53 | 0.53 | 0.53 | 0.53 | 0.5 | 0.53 |
| WSS quantile(0.95,$H_{\mathrm{E}}$) | 0.5 | 0.41 | 0.4 | 0.39 | 0.35 | 0.3 | 0.29 |
| WSS quantile(0.95, $D_{\mathrm{J}}$) | 0.49 | 0.32 | 0.12 | 0.11 | 0.06 | 0.06 | 0.07 |
| WSS quantile(0.95, $F_{\mathrm{ST}}$) | 0.64 | 0.51 | 0.25 | 0.25 | 0.15 | 0.16 | 0.18 |
| WSS quantile(0.95, $S$) | 14 | 18 | 16 | 18 | 20 | 30 | 27 |
| WSS quantile(0.95, $\theta_{\mathrm{W}}$) | 4.64 | 4.82 | 5.03 | 5.42 | 5.64 | 8.46 | 7.61 |
| WSS quantile(0.95, $\pi$) | 5.58 | 5.57 | 4.77 | 5.27 | 5.18 | 6.89 | 5.77 |
| WSS quantile(0.95, $D$) | 1.82 | 1.63 | 0.96 | 1 | 0.8 | 0.17 | 0.06 |
| WSS quantile(0.95, $D_{\mathrm{a}}$) | 0.38 | 0.23 | 0.08 | 0.08 | 0.04 | 0.04 | 0.05 |
| WSS $t_1$ quantile(0.95,$H_{\mathrm{E}}$) | 0.47 | 0.41 | 0.53 | 0.5 | 0.39 | 0.35 | 0.28 |
| WSS $t_2$ quantile(0.95,$H_{\mathrm{E}}$) | 0.53 | 0.37 | 0.42 | 0.4 | 0.36 | 0.3 | 0.32 |
| WSS $t_1$ quantile(0.95, $S$) | 4 | 15 | 10 | 11 | 16 | 13 | 13 |
| WSS $t_2$ quantile(0.95, $S$) | 13 | 15 | 14 | 17 | 15 | 28 | 24 |
| WSS $t_1$ quantile(0.95, $\theta_{\mathrm{W}}$) | 2.67 | 4.97 | 5.45 | 6 | 5.66 | 7.09 | 4.9 |
| WSS $t_2$ quantile(0.95, $\theta_{\mathrm{W}}$) | 4.6 | 4.97 | 4.95 | 5.63 | 5.3 | 8.44 | 7.95 |
| WSS $t_1$ quantile(0.95, $\pi$) | 2.67 | 5.2 | 5.5 | 6.17 | 5.62 | 7.17 | 4.66 |
| WSS $t_2$ quantile(0.95, $\pi$) | 5.6 | 4.67 | 4.8 | 5.3 | 5.09 | 7.01 | 6.56 |
| WSS $t_1$ quantile(0.95, $D$) | 3.19 | 1.45 | 1.63 | 1.63 | 1 | 1.09 | 1.01 |
| WSS $t_2$ quantile(0.95, $D$) | 2.09 | 1.14 | 1.05 | 1.06 | 1.05 | 0.3 | 0.38 |

**Table S3: Parameter posterior median and 95% credibility interval (in parenthesis) for each *A. mellifera* feral population.**

| Parameter | Avalon | Humboldt | Davis | Stanislaus | Stebbins | Riverside | Placerita |
|---|---|---|---|---|---|---|---|
| $P_R P_B$ | 1.53E-04<br>(1.59E-07 – 2.48E-01) | 8.35E-04<br>(1.55E-06 – 2.02E-01) | 1.26E-04<br>(1.17E-07 – 4.99E-02) | 2.39E-04<br>(4.32E-06 – 1.70E-01) | 9.70E-05<br>(1.14E-07 – 4.64E-02) | 8.24E-05<br>(1.18E-07 – 5.45E-01) | 6.19E-04<br>(4.15E-07 – 3.54E-01) |
| $P$ | 1.48E-05<br>(1.45E-06 – 1.04E-03) | 4.12E-05<br>(8.58E-06 – 3.08E-03) | 1.89E-05<br>(1.82E-06 – 3.99E-04) | 3.26E-05<br>(2.16E-06 – 3.49E-04) | 2.61E-05<br>(2.95E-06 – 5.23E-04) | 3.26E-05<br>(3.32E-06 – 4.93E-01) | 9.39E-05<br>(4.90E-06 – 4.93E-01) |
| $\gamma$ | 1.77E-02<br>(1.12E-03 – 8.82E-01) | 2.67E-02<br>(1.22E-03 – 8.45E-01) | 2.14E-02<br>(1.12E-03 – 9.12E-01) | 2.29E-02<br>(1.05E-03 – 8.93E-01) | 2.35E-02<br>(1.04E-03 – 8.93E-01) | 3.20E-02<br>(1.03E-03 – 8.93E-01) | 2.67E-02<br>(1.15E-03 – 8.92E-01) |
| $\bar{s}$ | 9.72E-02<br>(4.31E-03 – 1.62E+00) | 3.07E-01<br>(3.22E-02 – 2.23E+00) | 1.75E-01<br>(1.95E-03 – 1.50E+00) | 2.30E-01<br>(1.95E-03 – 1.30E+00) | 1.58E-01<br>(1.31E-03 – 1.51E+00) | 2.19E-01<br>(1.46E-03 – 2.25E+00) | 3.31E-01<br>(6.17E-03 – 1.98E+00) |
| $\theta_b$ | 3.72E-01<br>(1.72E-04 – 5.66E+03) | 1.73E+00<br>(1.22E-03 – 1.56E+04) | 2.90E+00<br>(1.70E-02 – 1.32E+03) | 3.46E+00<br>(2.66E-02 – 1.37E+03) | 3.07E+00<br>(9.20E-03 – 4.61E+02) | 2.90E+00<br>(4.41E-03 – 2.04E+04) | 6.22E+00<br>(1.30E-02 – 1.70E+04) |
| $L$ | 1.01E-03<br>(1.14E-09 – 9.64E-01) | 1.59E-04<br>(1.42E-16 – 9.27E-01) | 2.06E-01<br>(5.32E-08 – 8.29E-01) | 2.65E-01<br>(1.22E-06 – 8.78E-01) | 2.16E-04<br>(2.00E-16 – 4.62E-01) | 9.43E-03<br>(1.47E-08 – 8.71E-01) | 2.59E-02<br>(2.01E-16 – 7.56E-01) |
| $\mu$ | 5.67E-08<br>(3.45E-09 – 1.65E-07) | 8.05E-08<br>(9.75E-09 – 1.66E-07) | 5.03E-08<br>(4.04E-09 – 1.57E-07) | 2.56E-08<br>(3.06E-09 – 1.45E-07) | 7.12E-08<br>(4.86E-09 – 1.66E-07) | 2.98E-08<br>(2.71E-09 – 1.65E-07) | 1.11E-07<br>(1.23E-08 – 1.67E-07) |
| $r$ | 3.72E-08<br>(1.02E-08 – 6.14E-06) | 5.19E-08<br>(1.02E-08 – 2.06E-06) | 2.71E-08<br>(1.02E-08 – 2.01E-06) | 3.19E-08<br>(1.04E-08 – 1.06E-06) | 2.28E-08<br>(1.02E-08 – 8.54E-07) | 2.10E-08<br>(1.02E-08 – 1.06E-06) | 3.54E-08<br>(1.02E-08 – 3.64E-06) |
| $N$ | 8.10E+01<br>(1.70E+01 – 5.36E+03) | 4.90E+01<br>(1.00E+01 – 7.71E+03) | 1.02E+03<br>(2.42E+02 – 6.78E+03) | 1.80E+03<br>(3.67E+02 – 8.00E+03) | 3.42E+02<br>(1.58E+02 – 3.09E+03) | 5.19E+02<br>(6.30E+01 – 8.72E+03) | 1.39E+02<br>(1.40E+01 – 4.69E+03) |
| $N_e$ | 8.30E+01<br>(2.21E+01 – 9.59E+02) | 4.42E+01<br>(4.55E+00 – 5.76E+02) | 1.04E+03<br>(2.03E+02 – 7.24E+03) | 1.96E+03<br>(4.19E+02 – 8.70E+03) | 3.80E+02<br>(1.76E+02 – 3.90E+03) | 5.69E+02<br>(5.53E+01 – 8.17E+03) | 1.46E+02<br>(4.57E+00 – 1.15E+03) |
| $N_e/N$ | 9.07E-01<br>(1.15E-03 – 1.13E+00) | 1.05E+00<br>(8.75E-04 – 1.17E+00) | 1.07E+00<br>(5.63E-01 – 1.12E+00) | 1.06E+00<br>(5.26E-01 – 1.12E+00) | 1.09E+00<br>(2.28E-01 – 1.13E+00) | 1.04E+00<br>(2.78E-01 – 1.13E+00) | 1.06E+00<br>(2.44E-01 – 1.14E+00) |
| $F_{ST}$-$N_e$ | 8.00E+01<br>(0.00E+00 – 7.80E+02) | 1.13E+02<br>(5.30E+00 – 5.88E+17) | 3.62E+01<br>(8.63E+00 – 3.59E+02) | 2.28E+01<br>(6.00E+00 – 3.42E+02) | 3.60E+01<br>(5.63E+00 – 2.25E+02) | 1.47E+01<br>(1.79E+00 – 1.20E+02) | 6.15E+01<br>(4.46E+00 – 6.12E+02) |

Estimates are in the original scale.

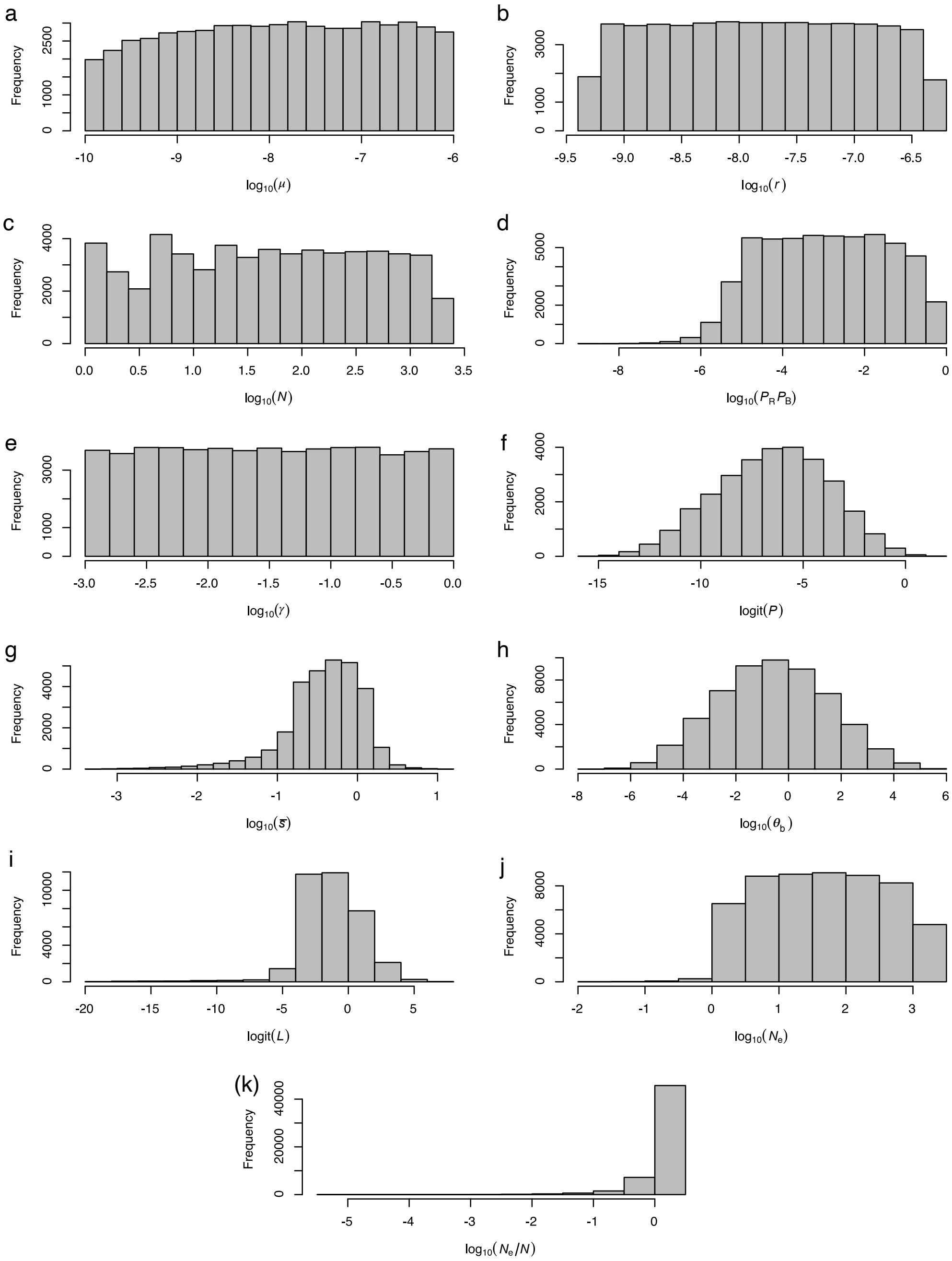


**Figure S1: Prior distribution of the simulation parameters, latent variables informative about the adaptive history and demography inferred with the ABC-RF.** (a) the per-site mutation rate $\mu$, (b) per base recombination rate per generation $r$, (c) the population size $N$, (d) the probability of beneficial mutation $P_R P_B$, (e) mean of the gamma distribution $\gamma$, (f) the proportion of strongly selected mutations $P$, (g) average selection coefficients of mutations under strong selection $\bar{s}$, (h) the scale mutation rate of beneficial mutations $\theta_{\mathrm{b}}$, (i) the average substitution load $L$, (j) the effective population size $N_{\mathrm{e}}$, (k) the distance between the effective size and population size expressed in the ratio $N_{\mathrm{e}}/N$.

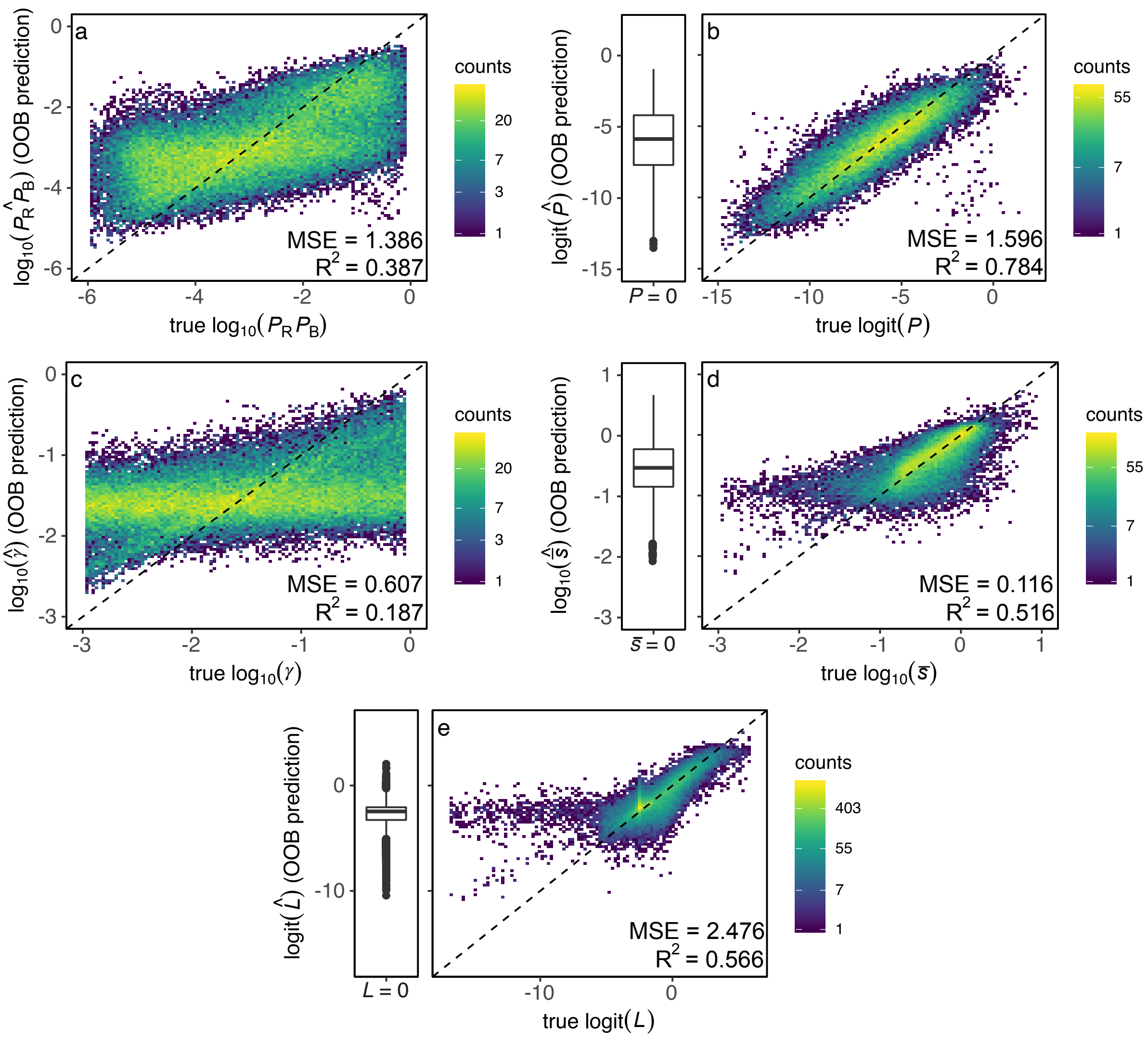


**Figure S2: Out-of-bag estimates of ABC-RF trained for the prediction of model parameters informative about the number of beneficial mutations and their strength, the latent variables associated with these two parameters, and variables informative about the adaptive dynamic.** (a) probability of a beneficial mutation to arise $P_R P_B$; (b) number of mutations under strong selection $P$; (c) mean of the gamma distribution $\gamma$; (d) mean of selection coefficients of mutations under strong selection $\bar{s}$; and (e) mean substitution load $L$. The bar plots on the left side of the scatterplots of $P$, $\bar{s}$, and $L$ correspond to the posterior estimates obtained for the simulations removed for training these parameters, as these simulations had zero as true values. MSE and $R^2$ estimates of b, d, and e corresponded to non-neutral simulations. See Table S2, Supplementary results, for neutral simulations MSE and bias.

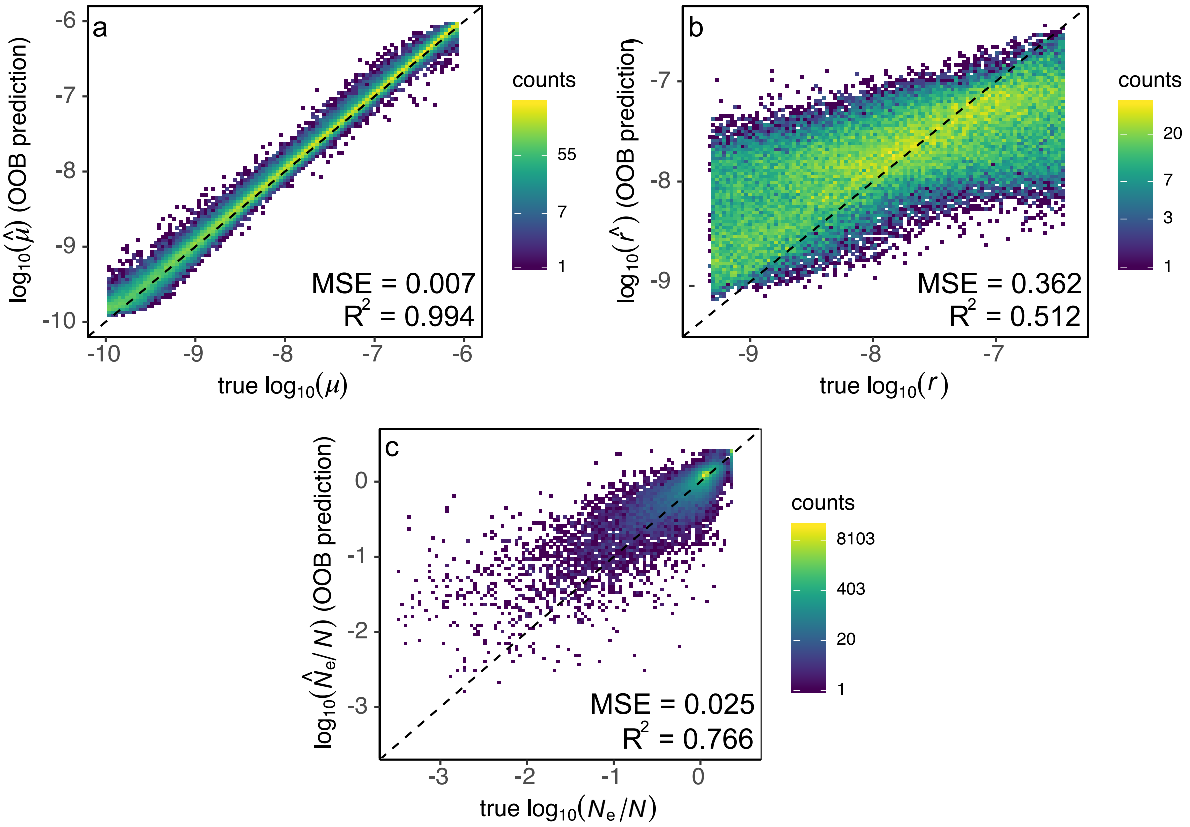


**Figure S3: Out-of-bag estimates of ABC-RF trained for predicting the model parameters and latent variables informative about demography.** (a) mutation rate per generation $\mu$; (b) per base recombination rate per generation $r$; and (c) the ratio between the effective size and the population census size $N_\mathrm{e}/N$.

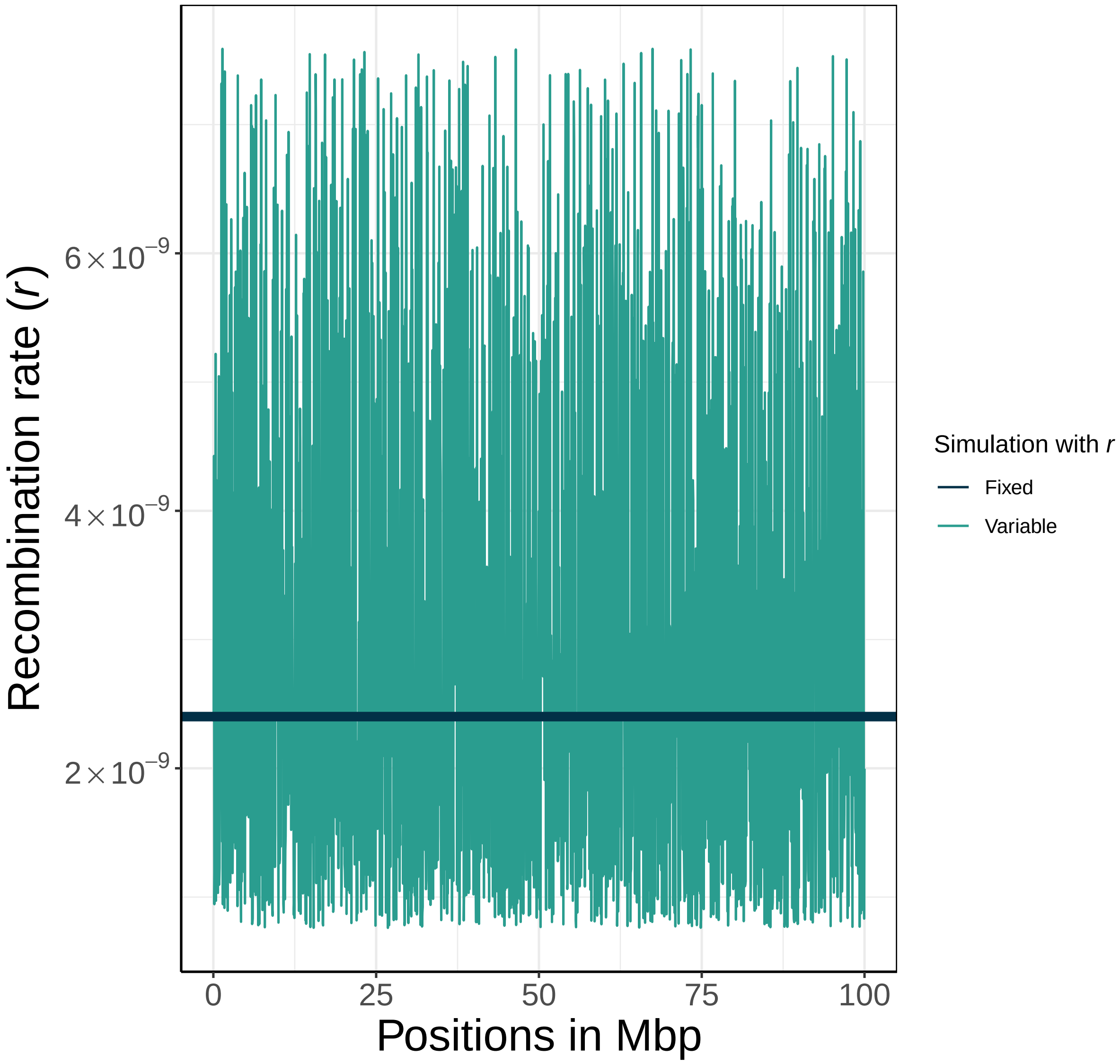


**Figure S4: How the recombination rate per base pair per generation $r$ varies across the genome.** Simulation for model training had constant $r$ (horizontal line in dark blue). Simulations used to test the ABC-RF robustness to heterogenous recombination rates could have different values of $r$ for the recombination map. This figure depicts one example of genome-wide $r$ for each type of simulation.

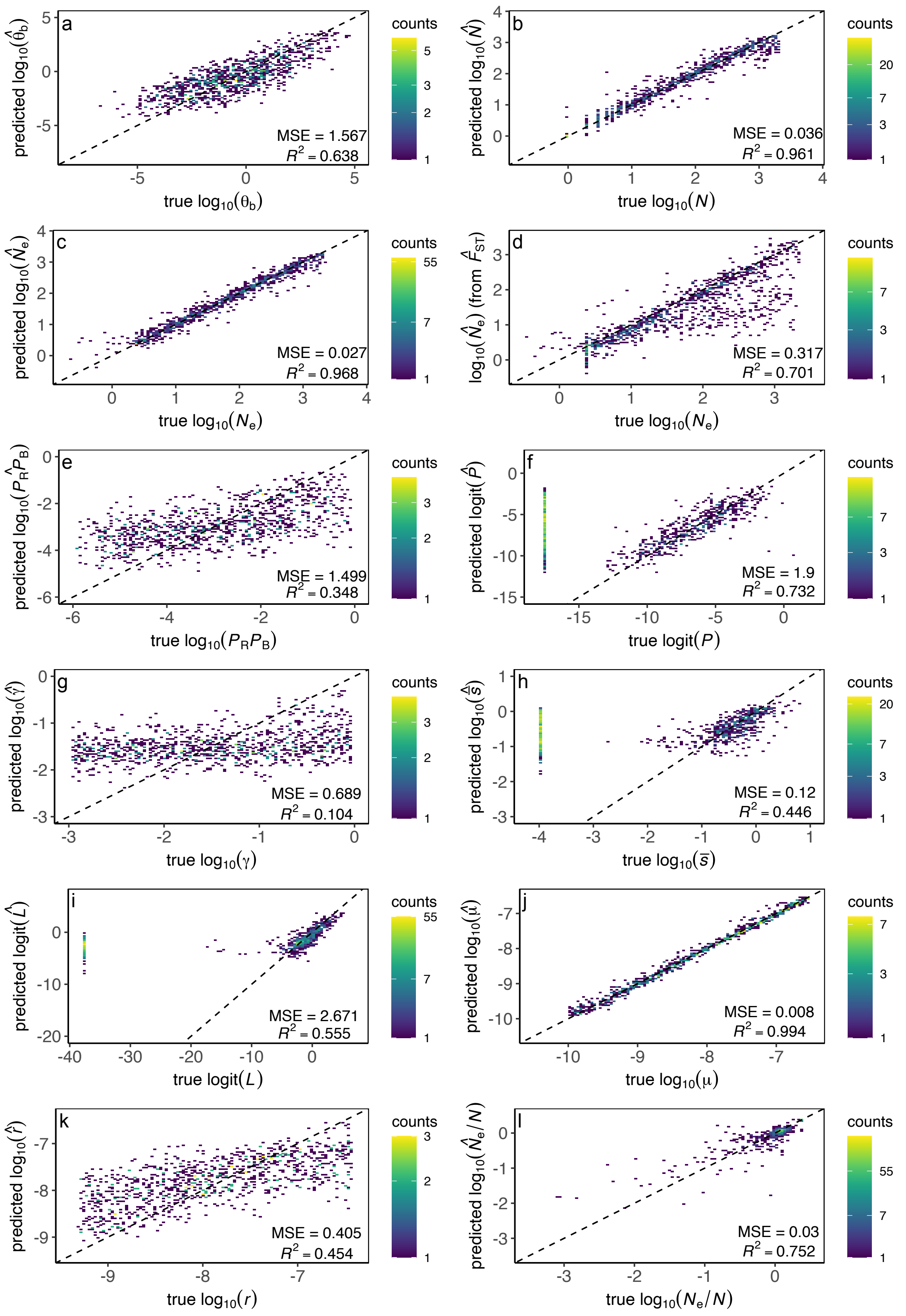


**Figure S5: Expected versus posterior estimates plots for PODs with variable genome-wide recombination rates.** (a) $\theta_\mathrm{b}$, (b) $N$, (c) $N_\mathrm{e}$, (d) $F_\mathrm{ST} - N_\mathrm{e}$, (e) $P_R P_B$, (f) $P$, (g) $\gamma$, (h) $\bar{s}$, (i) $L$, (j) $\mu$, (k) $r$, and (l) $N_\mathrm{e}/N$. MSE and $R^2$ estimates of f, h, and i corresponded to non-neutral PODs. See Table S2, Supplementary results, for neutral PODs MSE and bias.

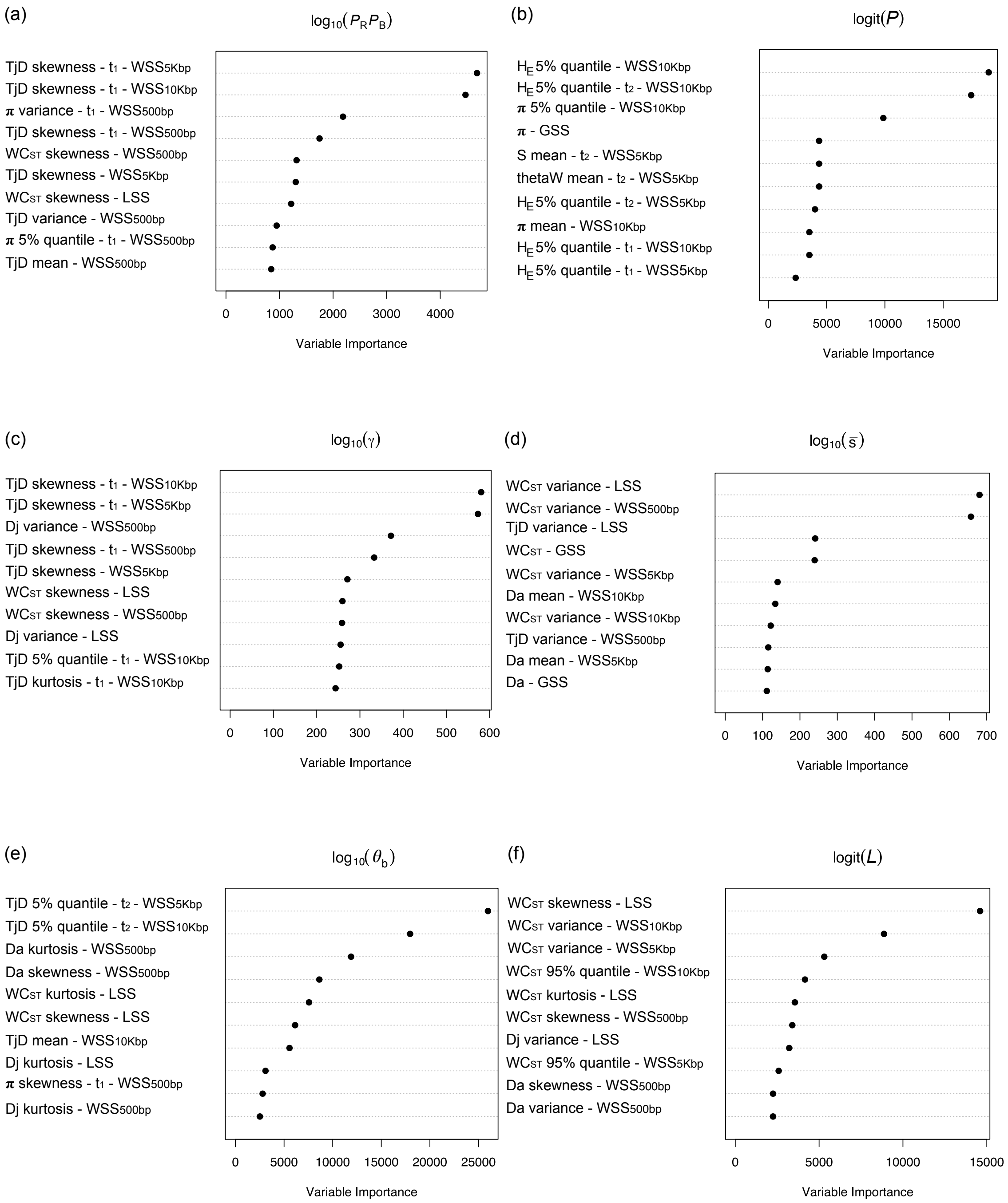


**Figure S6: Variable importance plots of the trained ABC-RF for parameters informative about the adaptive history.** (a) The probability of beneficial mutation $P_R P_B$, (b) the proportion of strongly selected mutations $P$, (c) mean of the gamma distribution $\gamma$, (d) average selection coefficients of mutations under strong selection $\bar{s}$, (e) the scale mutation rate of selected mutations $\theta_{\mathrm{b}}$, and (f) the average substitution load $L$.

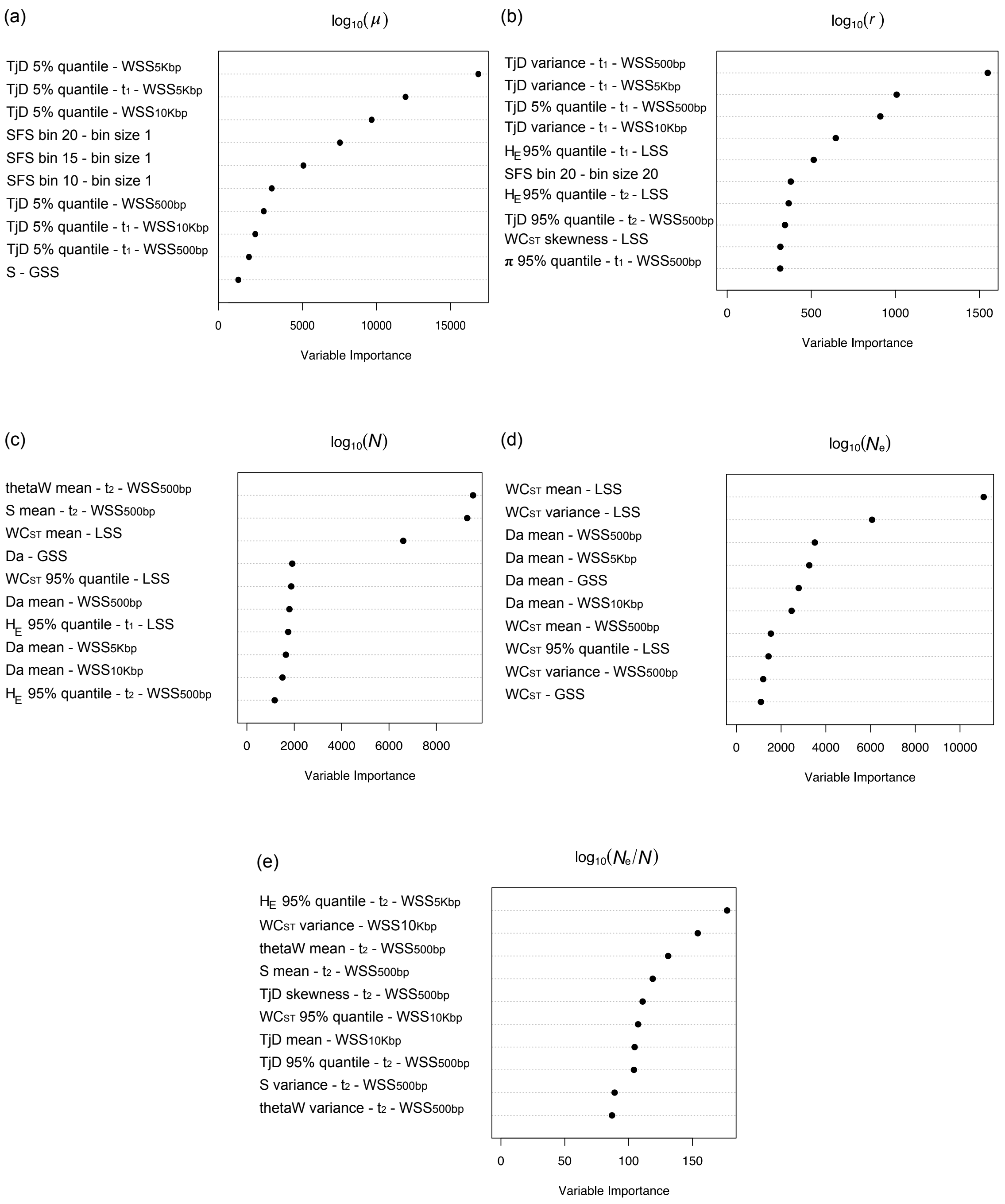


**Figure S7: Variable importance plots of the trained ABC-RF for parameters informative about the adaptive history.** (a) the per-site mutation rate $\mu$, (b) per base recombination rate per generation $r$, (c) the population size $N$, (d) the effective population size $N_\mathrm{e}$, (e) the distance between the effective size and population size expressed in the ratio $N_\mathrm{e}/N$.

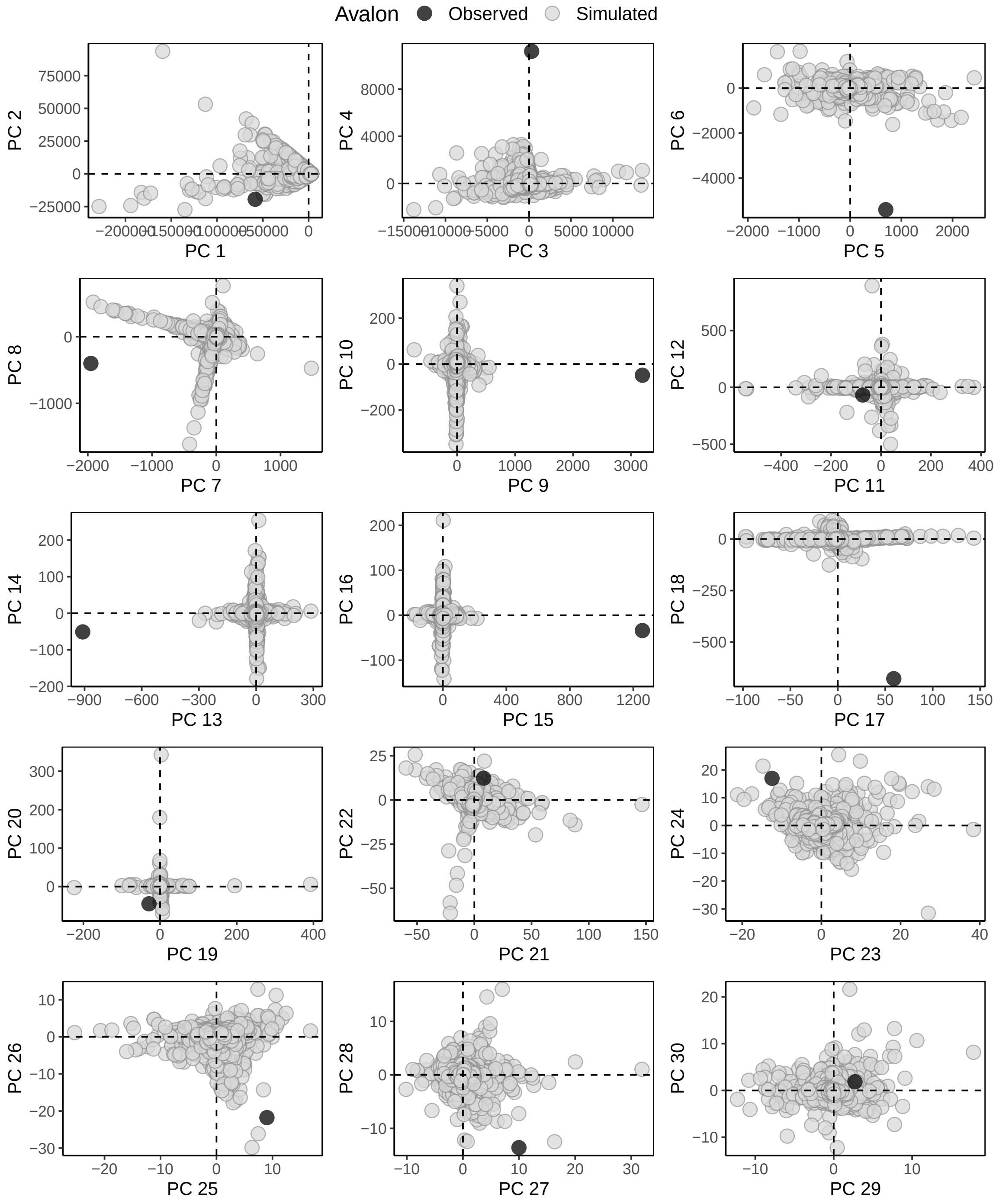


**Figure S8: Prior evaluation for the Avalon population.** Projection of the targeted population principal components, computed from the summary statistics, into the cloud of the simulated data points obtained with principal component analysis (PCA). We see the targeted data point (in black) into the simulated data cloud of points (in grey) if the model captured most of the targeted data features.

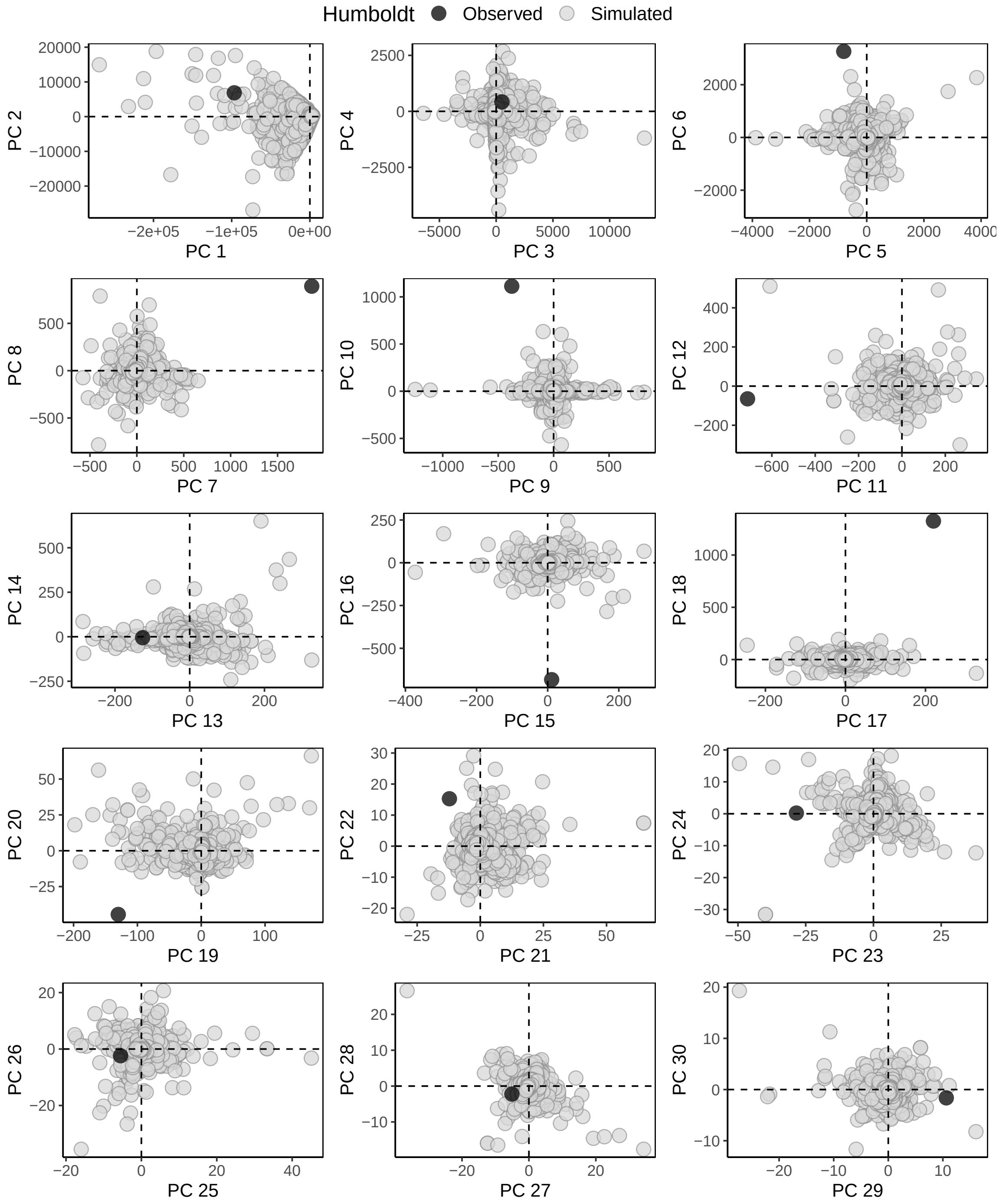


**Figure S9: Prior evaluation for the Humboldt population.** Projection of the targeted population principal components, computed from the summary statistics, into the cloud of the simulated data points obtained with principal component analysis (PCA). We see the targeted data point (in black) into the simulated data cloud of points (in grey) if the model captured most of the targeted data features.

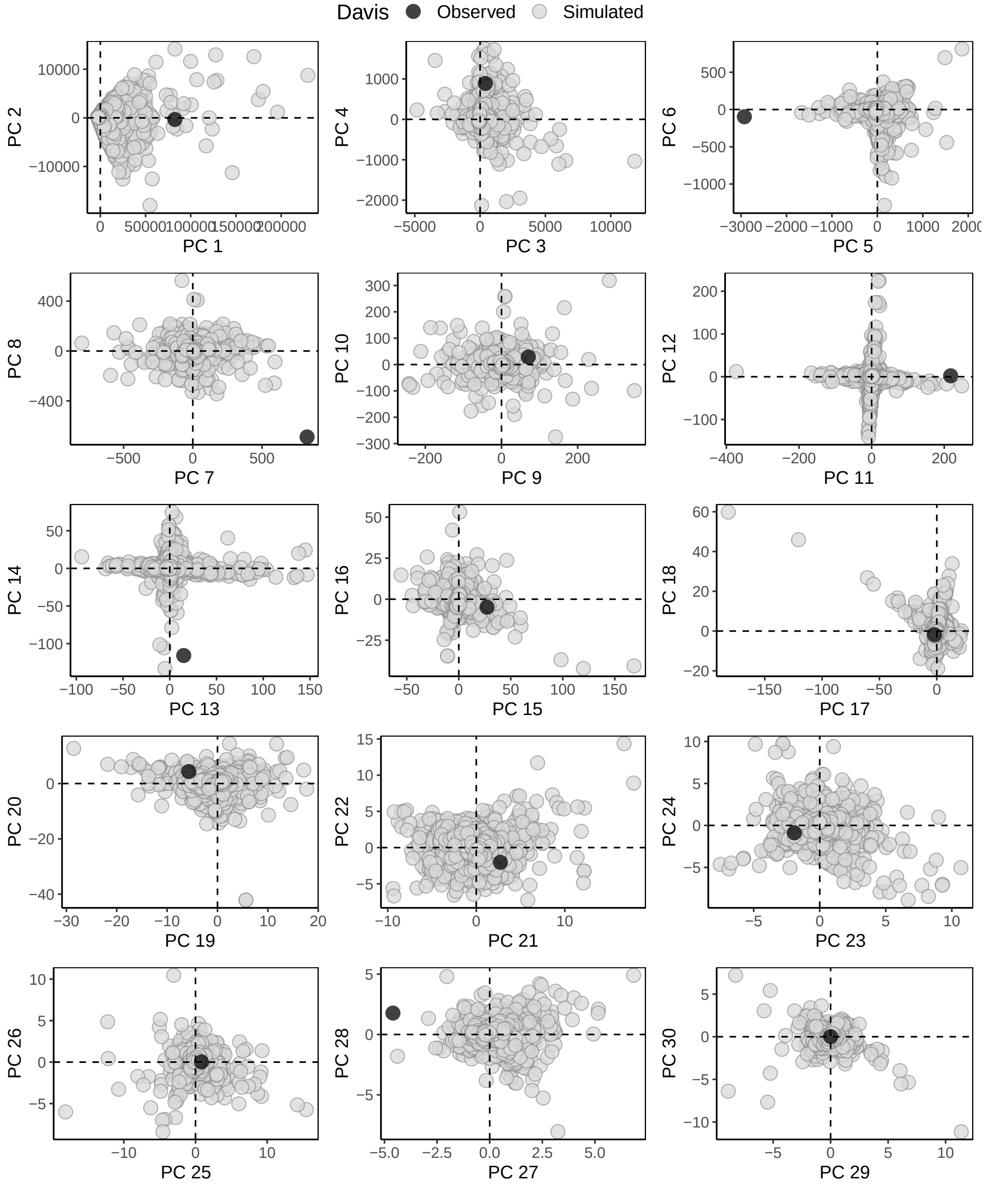


**Figure S10: Prior evaluation for the Davis population.** Projection of the targeted population principal components, computed from the summary statistics, into the cloud of the simulated data points obtained with principal component analysis (PCA). We see the targeted data point (in black) into the simulated data cloud of points (in grey) if the model captured most of the targeted data features.

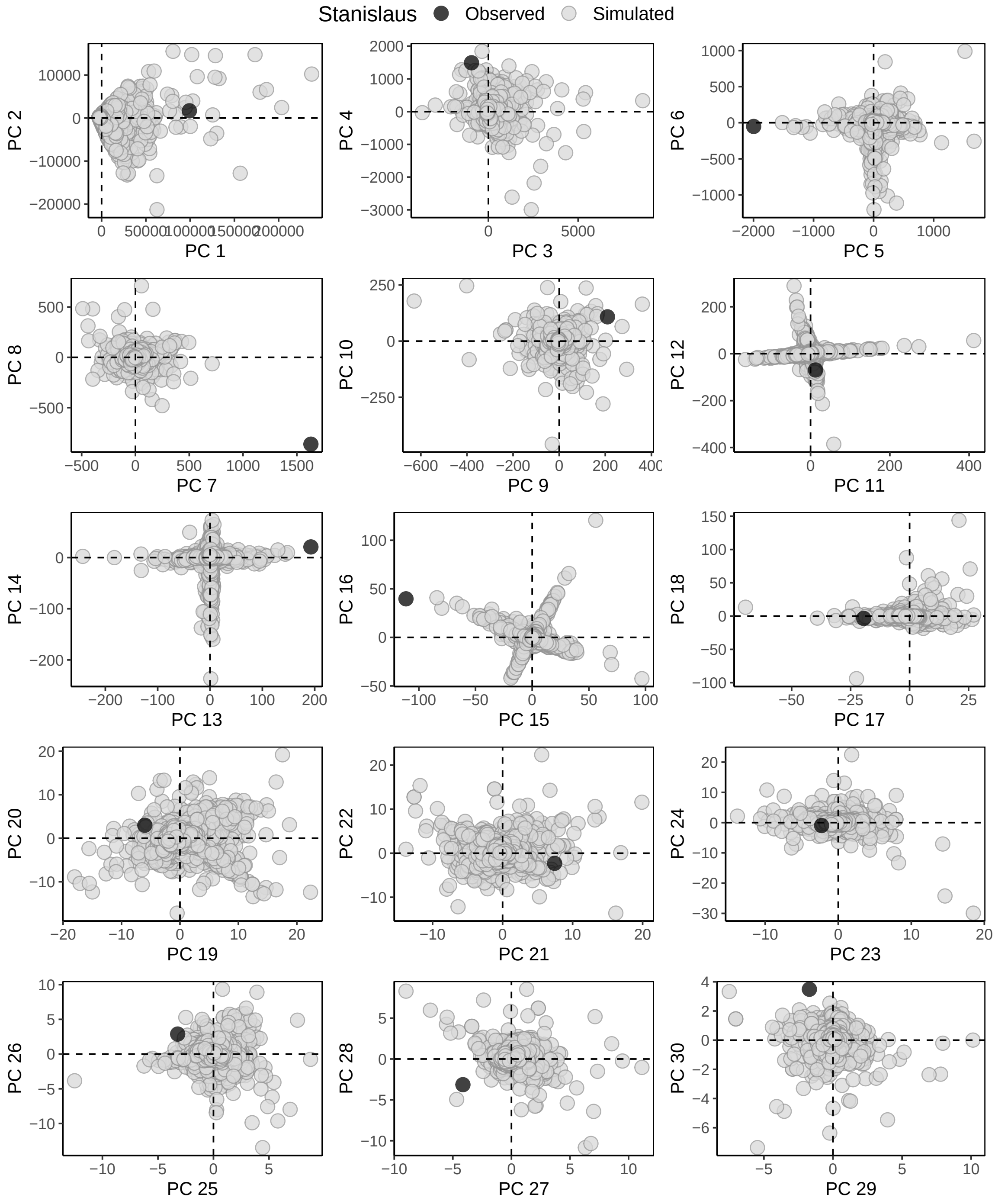


**Figure S11: Prior evaluation for the Stanislaus population.** Projection of the targeted population principal components, computed from the summary statistics, into the cloud of the simulated data points obtained with principal component analysis (PCA). We see the targeted data point (in black) into the simulated data cloud of points (in grey) if the model captured most of the targeted data features.

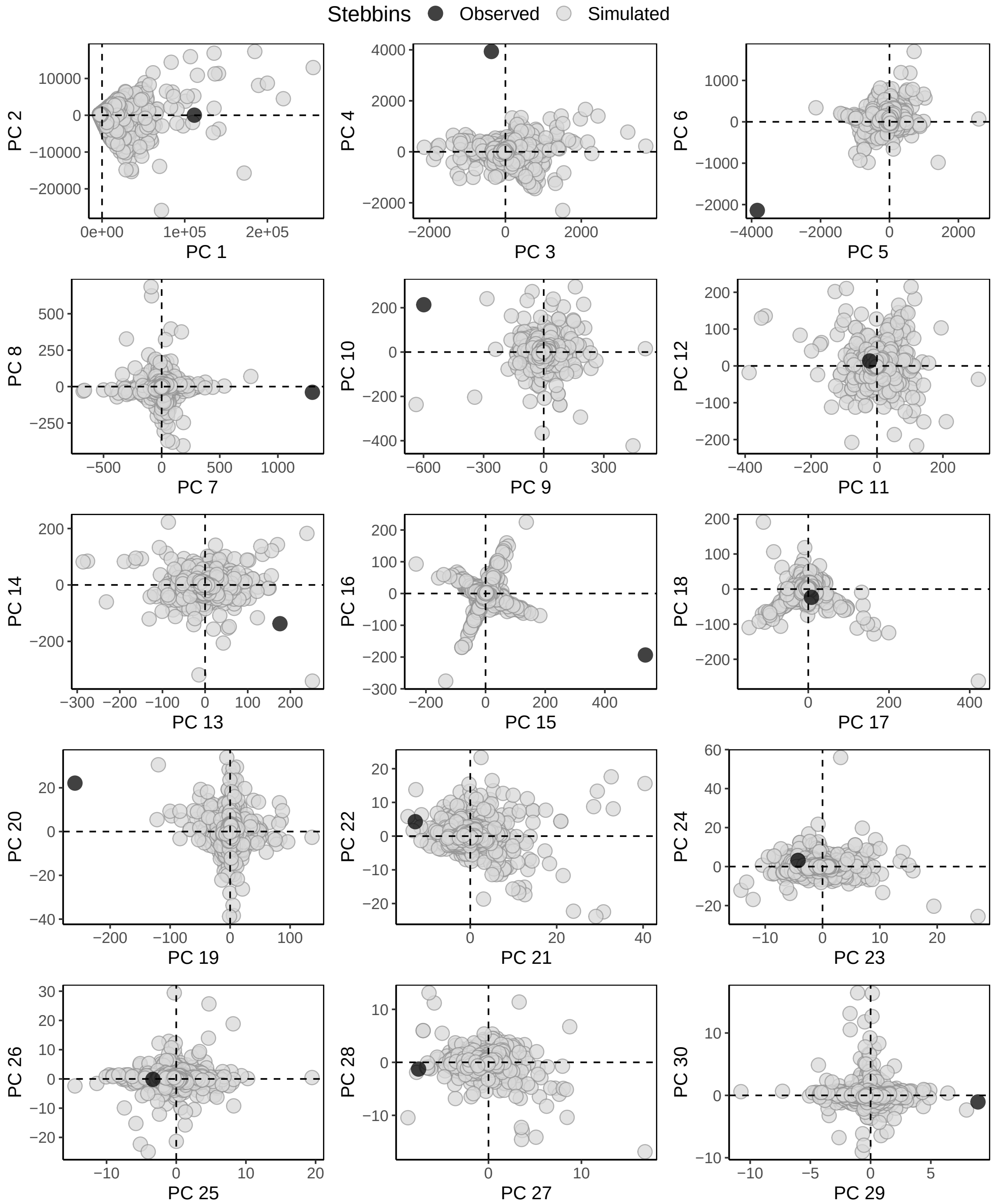


**Figure S12: Prior evaluation for the Stebbins population.** Projection of the targeted population principal components, computed from the summary statistics, into the cloud of the simulated data points obtained with principal component analysis (PCA). We see the targeted data point (in black) into the simulated data cloud of points (in grey) if the model captured most of the targeted data features.

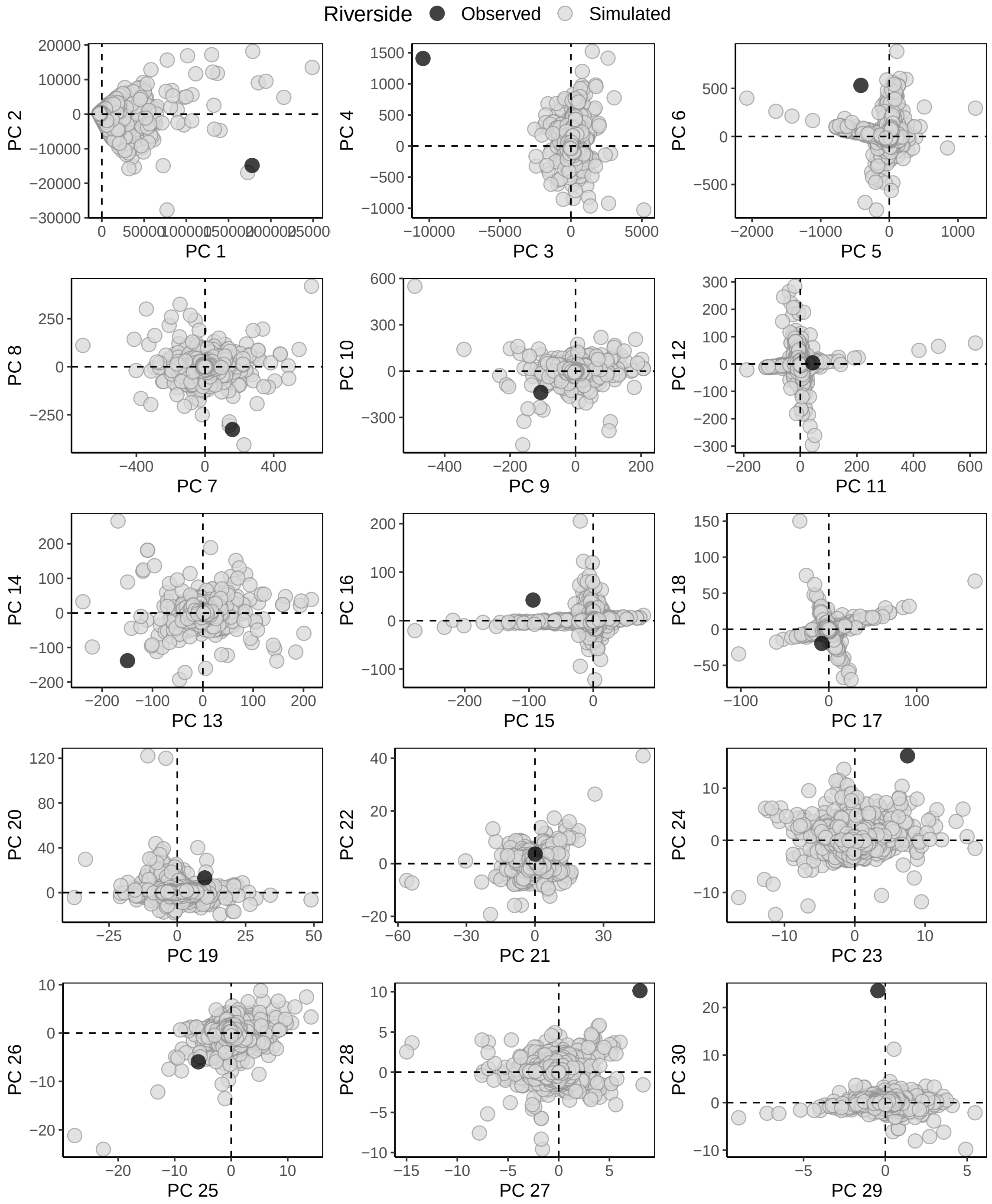


**Figure S13: Prior evaluation for the Riverside population.** Projection of the targeted population principal components, computed from the summary statistics, into the cloud of the simulated data points obtained with principal component analysis (PCA). We see the targeted data point (in black) into the simulated data cloud of points (in grey) if the model captured most of the targeted data features.

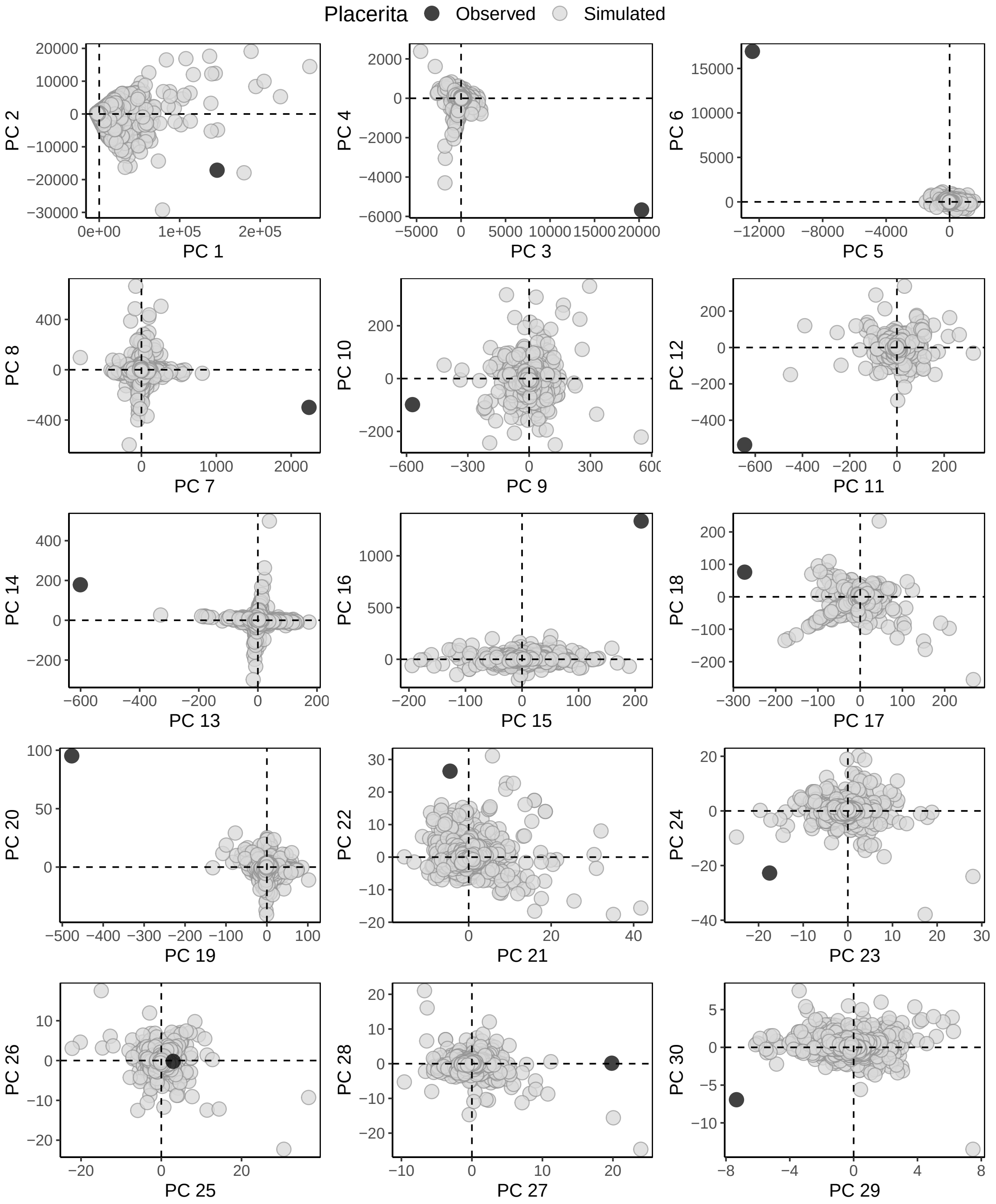


**Figure S14: Prior evaluation for the Placerita population.** Projection of the targeted population principal components, computed from the summary statistics, into the cloud of the simulated data points obtained with principal component analysis (PCA). We see the targeted data point (in black) into the simulated data cloud of points (in grey) if the model captured most of the targeted data features.

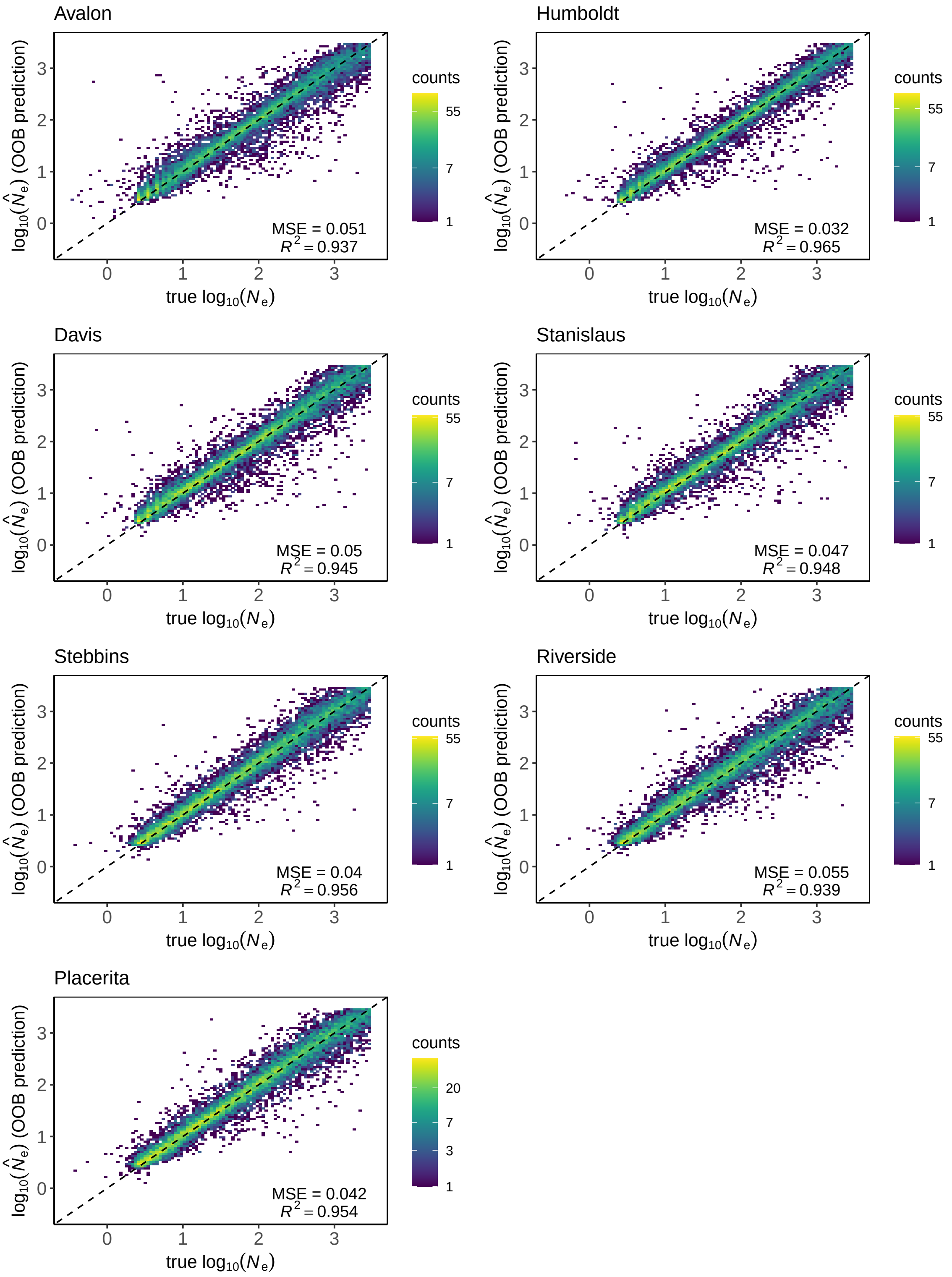


**Figure S15: Out-of-bag estimates of ABC-RF trained for predicting the population effective size $N_e$ for pairs of temporal samples of *A. mellifera* feral populations.**

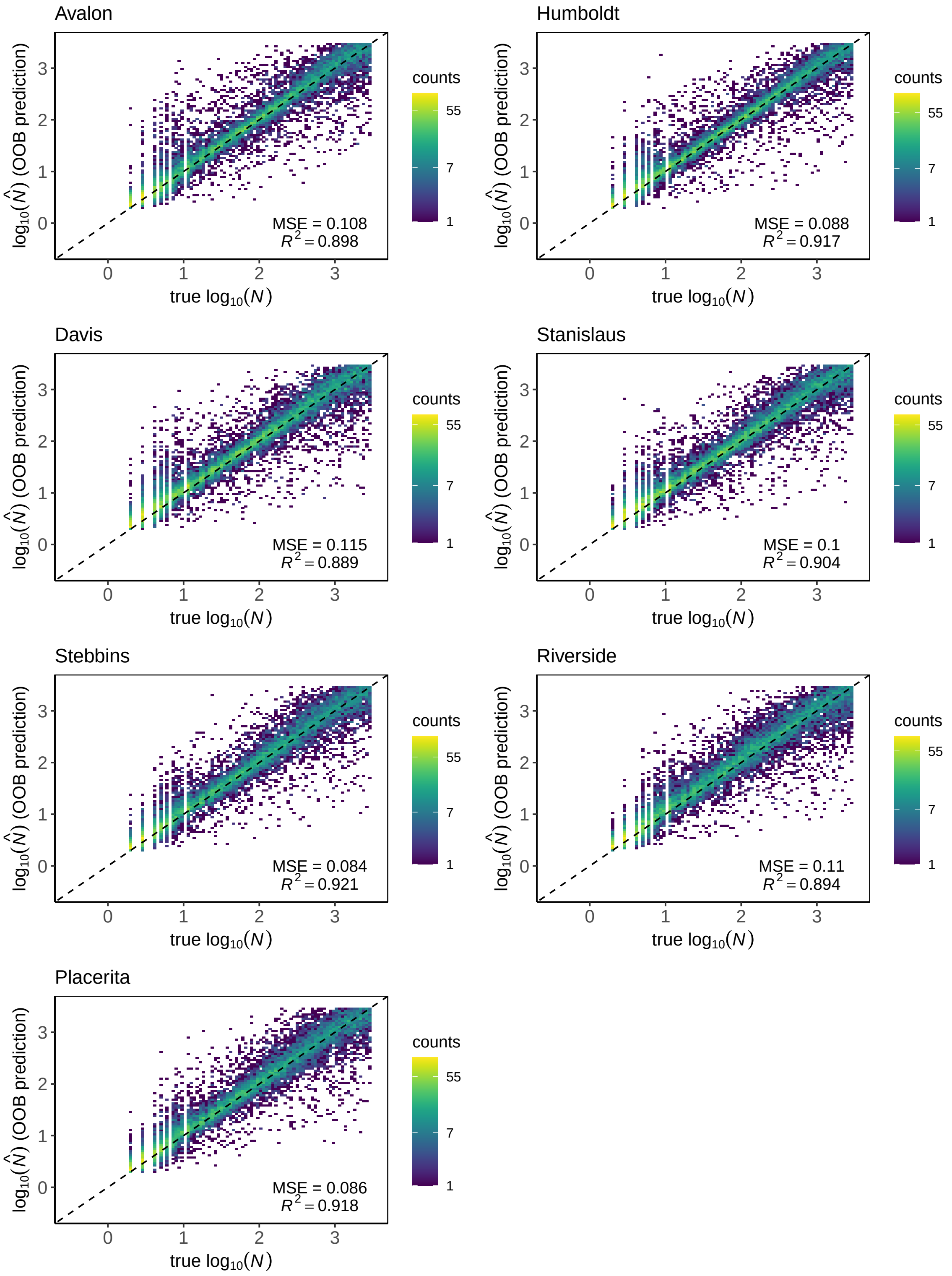


**Figure S16: Out-of-bag estimates of ABC-RF trained for predicting the population census size $N$ for pairs of temporal samples of *A. mellifera* feral populations.**

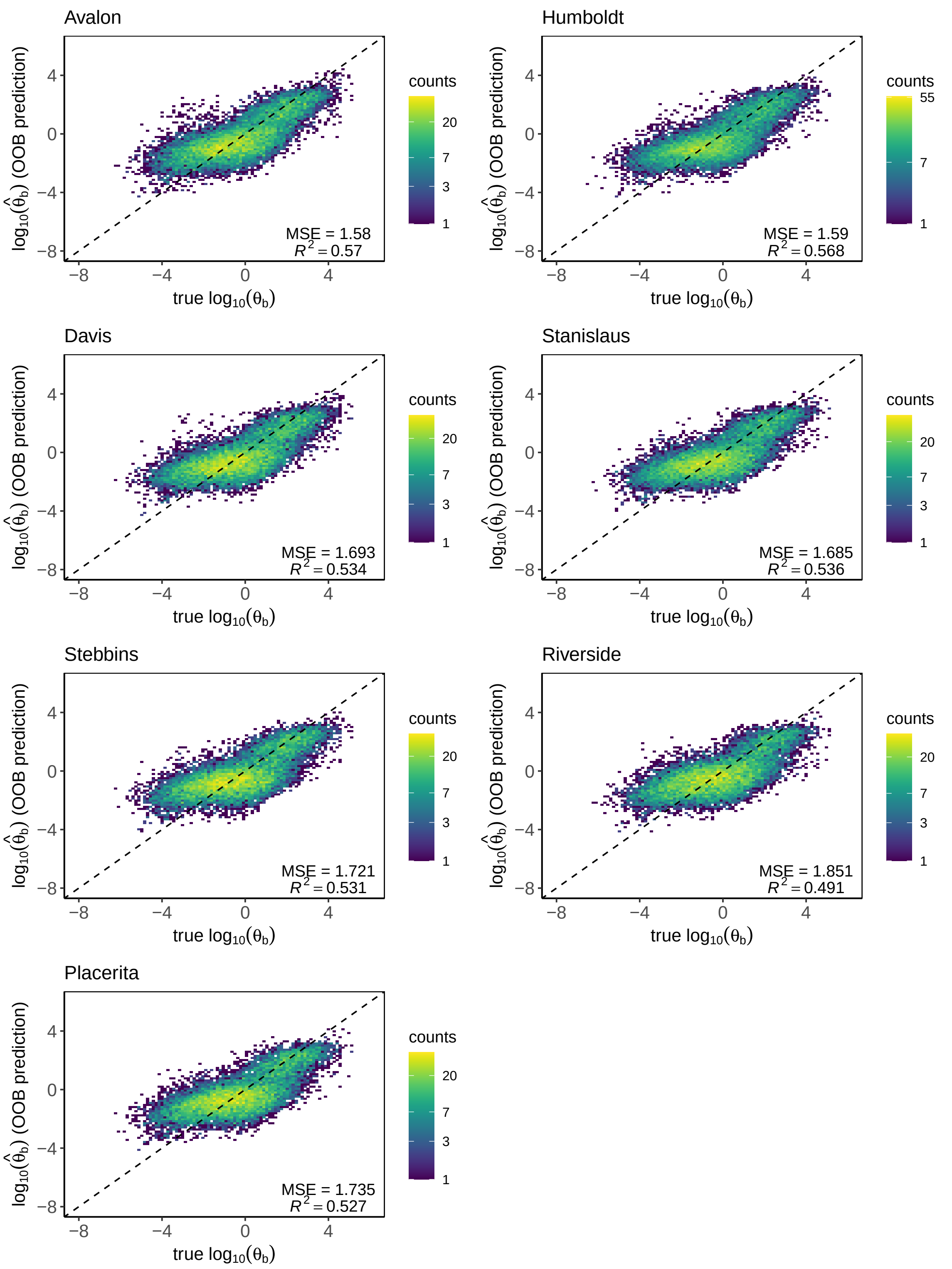


**Figure S17: Out-of-bag estimates of ABC-RF trained for predicting the scaled mutation rate of beneficial mutations $\theta_b$ for pairs of temporal samples of *A. mellifera* feral populations.**

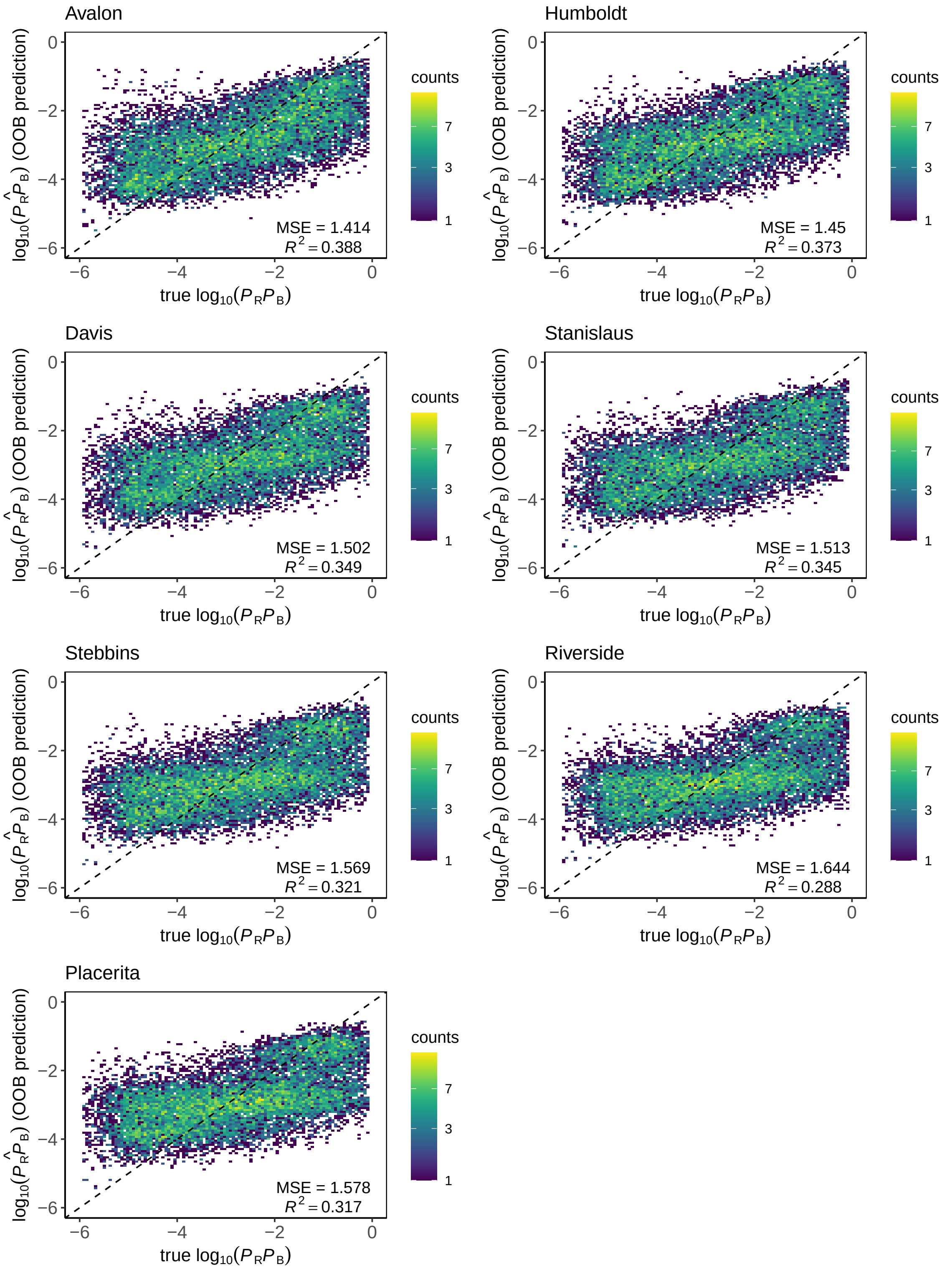


**Figure S18: Out-of-bag estimates of ABC-RF trained for predicting the probability of beneficial mutation $P_R P_B$ for pairs of temporal samples of *A. mellifera* feral populations.**

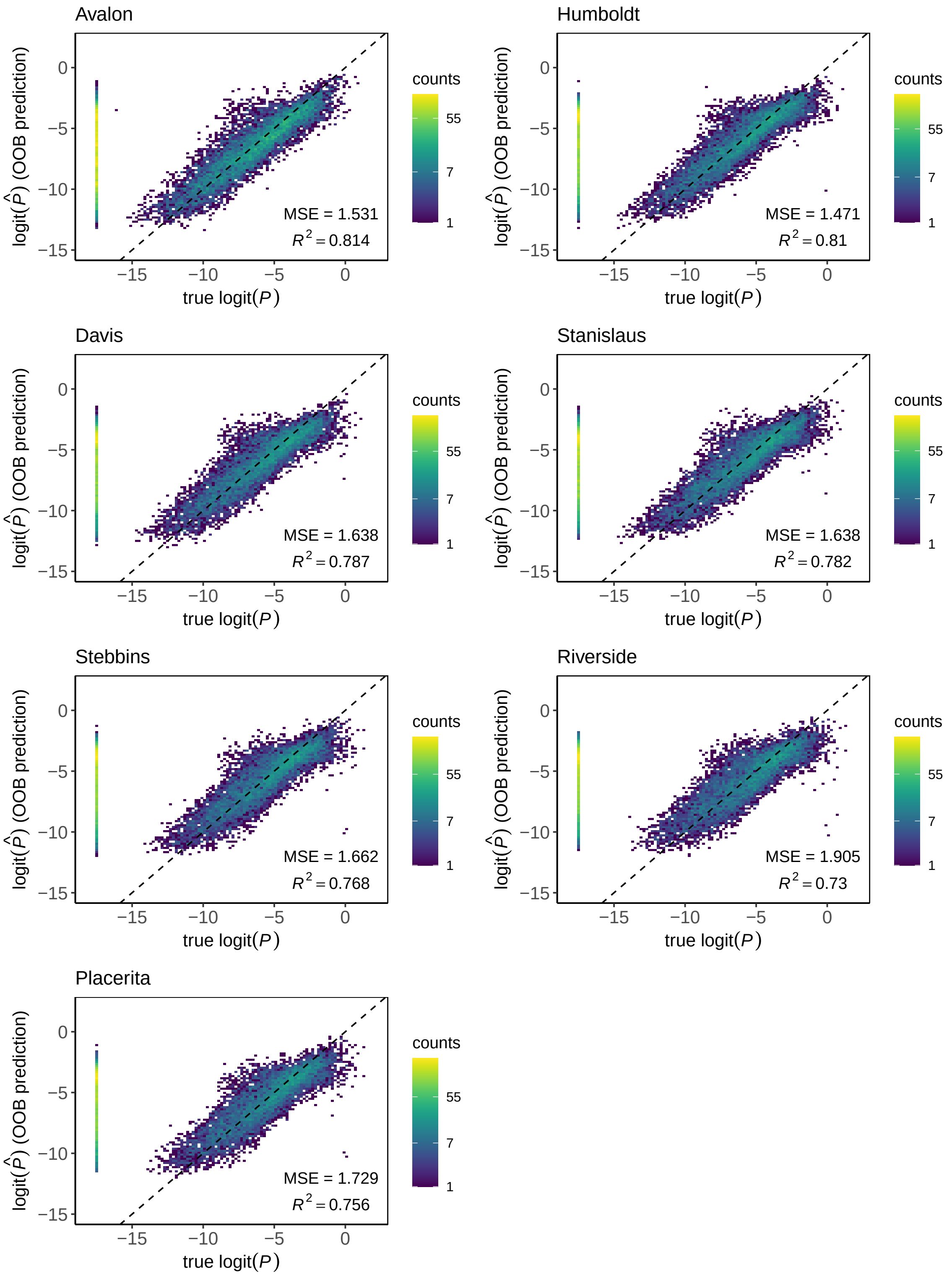


**Figure S19: Out-of-bag estimates of ABC-RF trained for predicting the proportion of strongly selected mutations $P$ for pairs temporal samples of *A. mellifera* feral populations.** Stacked points on the far left of each scatter-plot correspond to posterior estimates of neutral simulations, where $P = 0$. MSE and $R^2$ estimates corresponded to non-neutral simulations. See Table S2, Supplementary results, for neutral simulations MSE and bias.

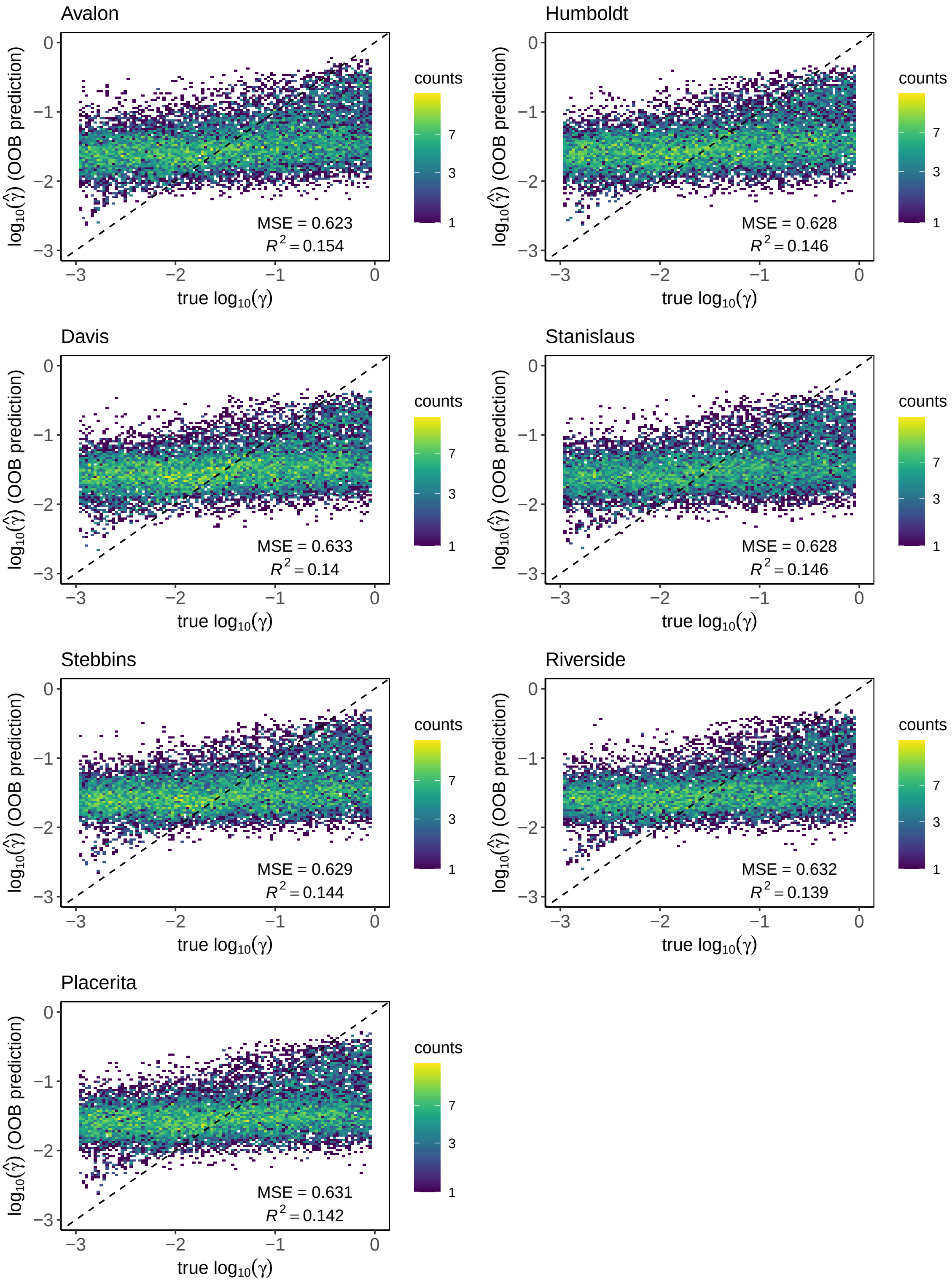


**Figure S20: Out-of-bag estimates of ABC-RF trained for predicting the mean of the gamma distribution $\gamma$ for pairs of temporal samples of *A. mellifera* feral populations.**

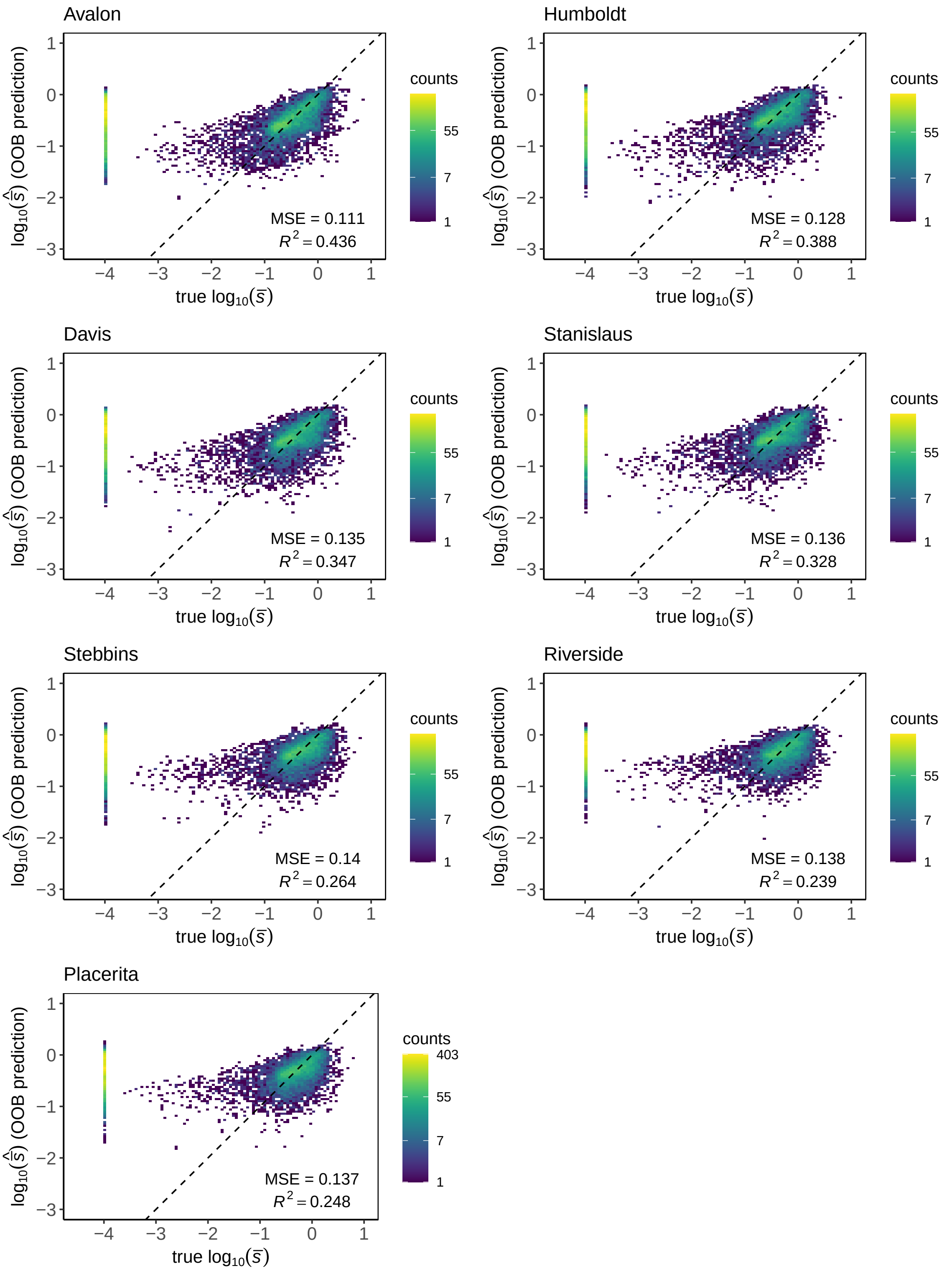


**Figure S21: Out-of-bag estimates of ABC-RF trained for predicting the average selection coefficients of mutations under strong selection $\bar{s}$ for pairs of temporal samples of *A. mellifera* feral populations.** Stacked points on the far left of each scatter-plot correspond to posterior estimates of neutral simulations, where $\bar{s} = 0$. MSE and $R^2$ estimates corresponded to non-neutral simulations. See Table S2, Supplementary results, for neutral simulations MSE and bias.

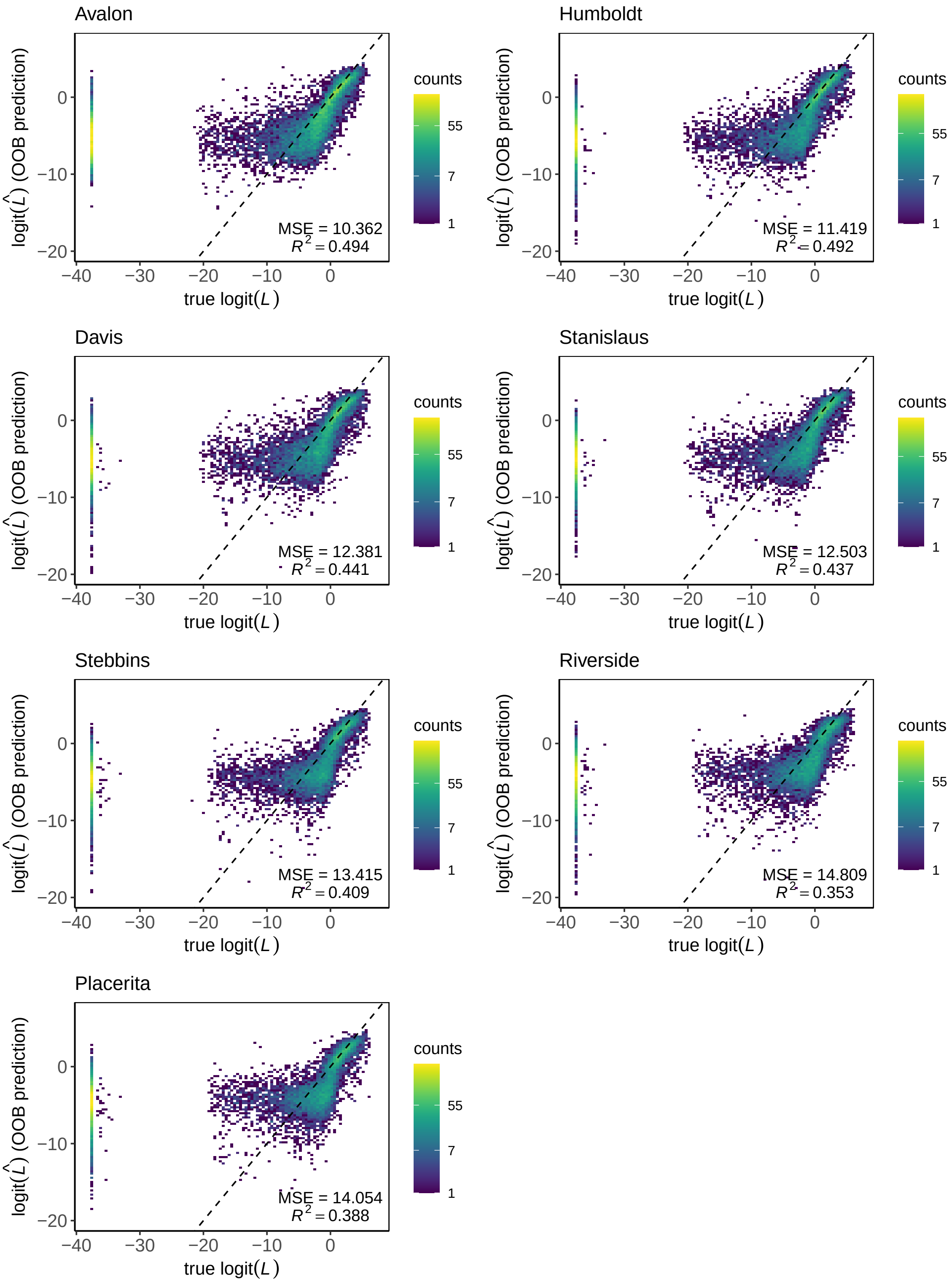


**Figure S22: Out-of-bag estimates of ABC-RF trained for predicting the mean substitution load $L$ for pairs of temporal samples *A. mellifera* feral populations.** Stacked points on the far left of each scatter-plot correspond to posterior estimates of neutral simulations, where $L = 0$. MSE and $R^2$ estimates corresponded to non-neutral simulations. See Table S2, Supplementary results, for neutral simulations MSE and bias.

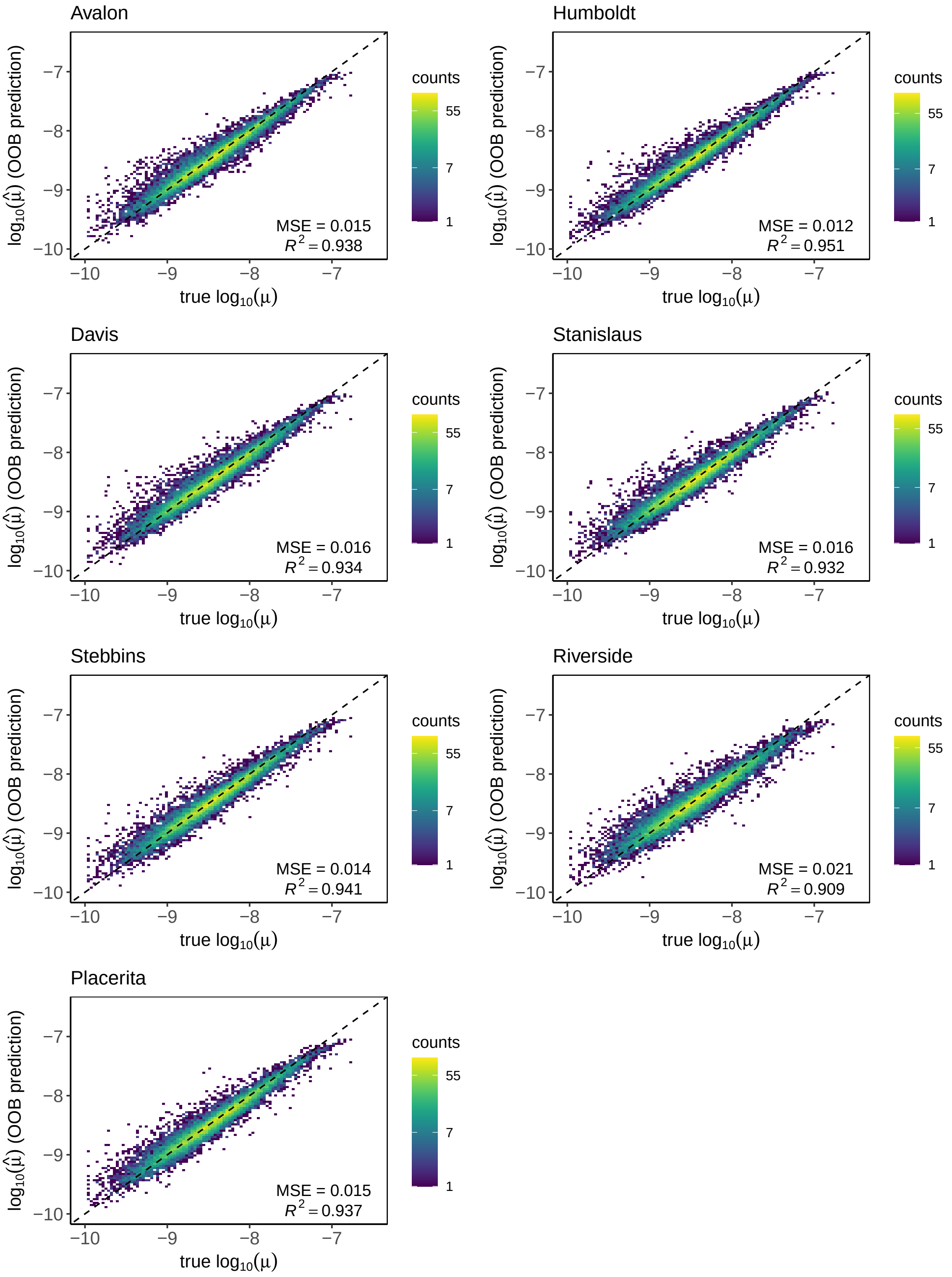


**Figure S23: Out-of-bag estimates of ABC-RF trained for the mutation rate per generation $\mu$ for pairs of temporal samples of *A. mellifera* feral populations.**

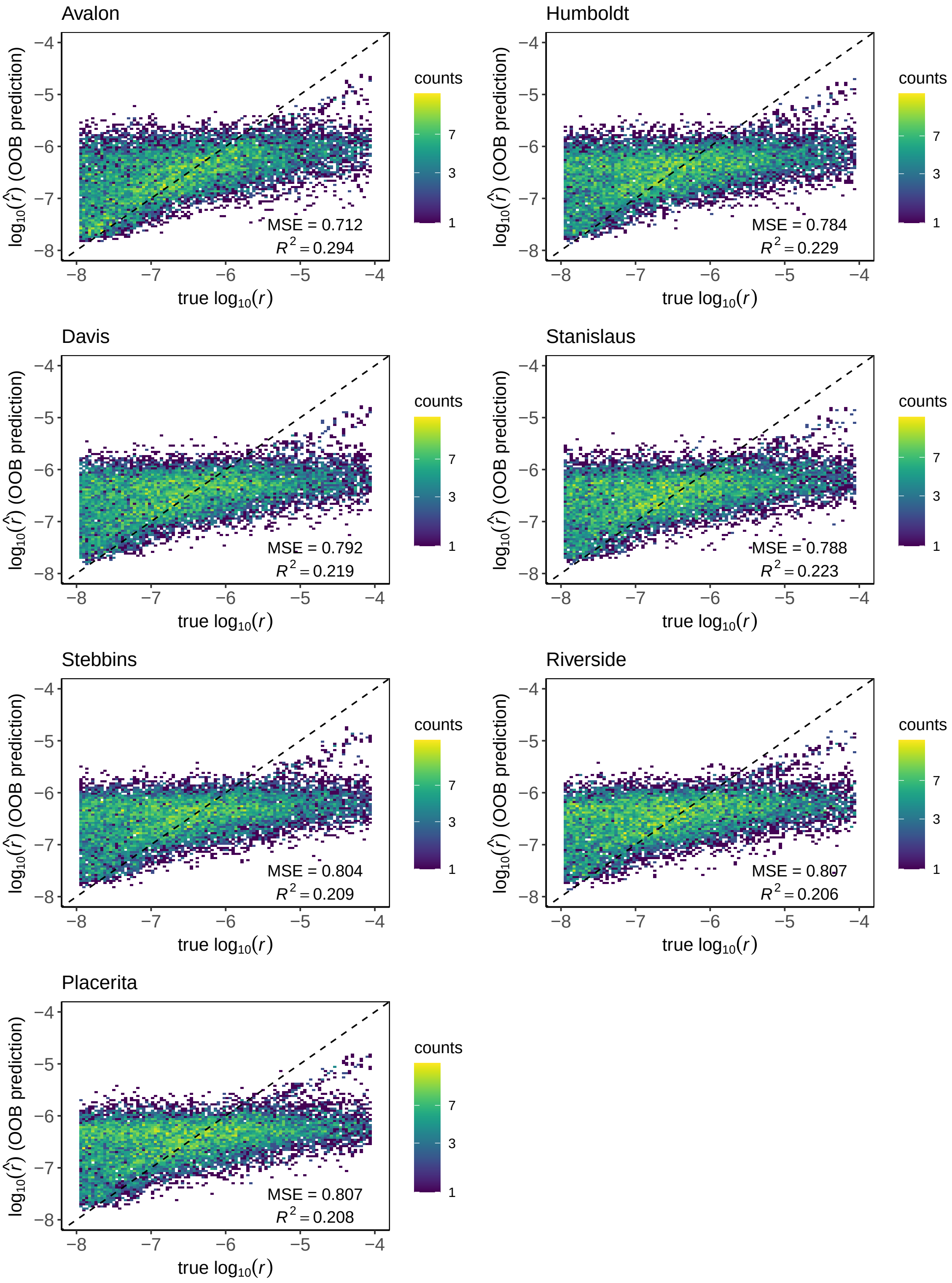


**Figure S24: Out-of-bag estimates of ABC-RF trained for predicting the per base recombination rate per generation $r$ for pairs of temporal samples of *A. mellifera* feral populations.**

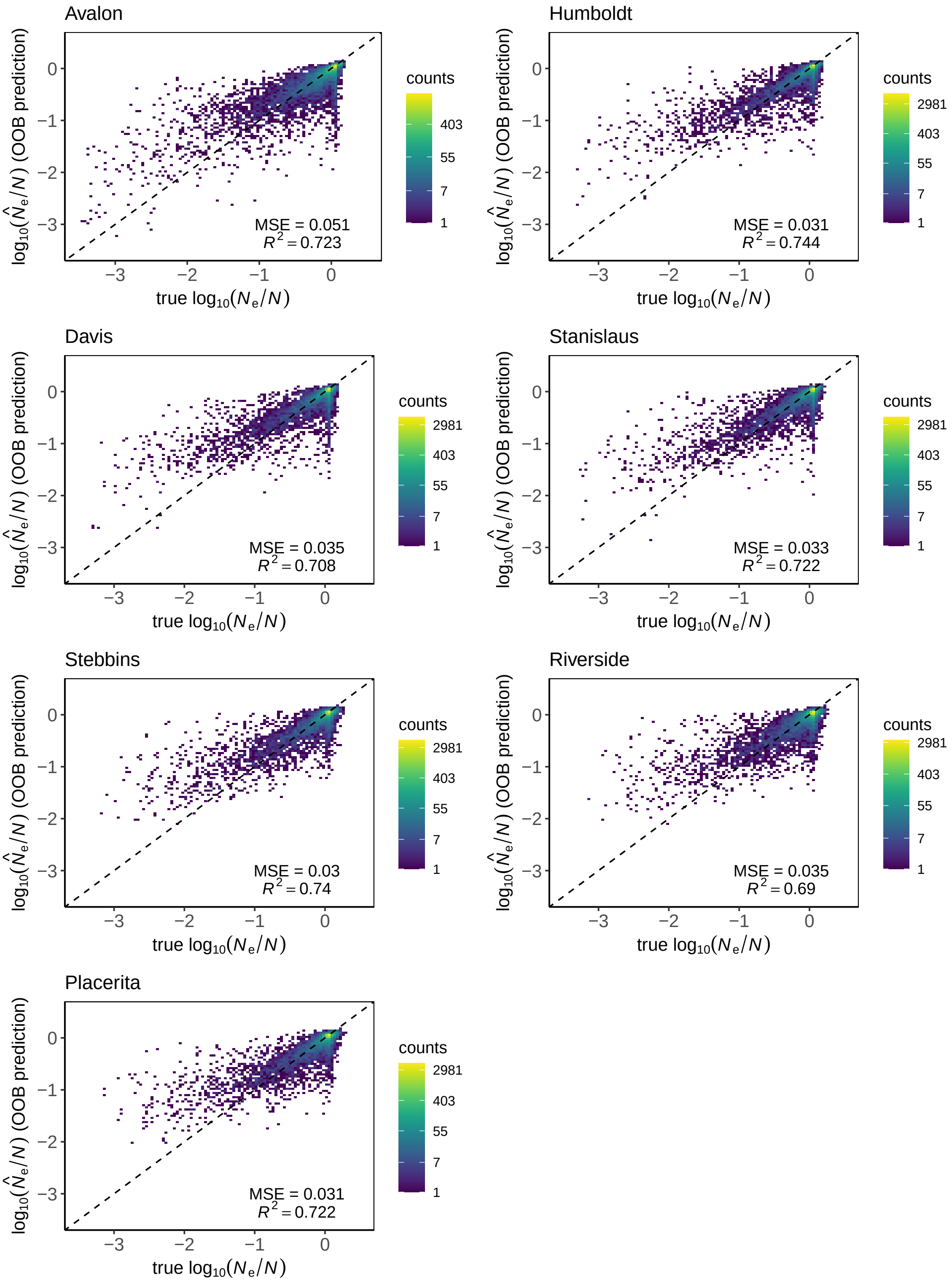


**Figure S25: Out-of-bag estimates of ABC-RF trained for predicting the ratio between the effective size and the population census size for pairs of temporal samples of *A. mellifera* feral populations.**

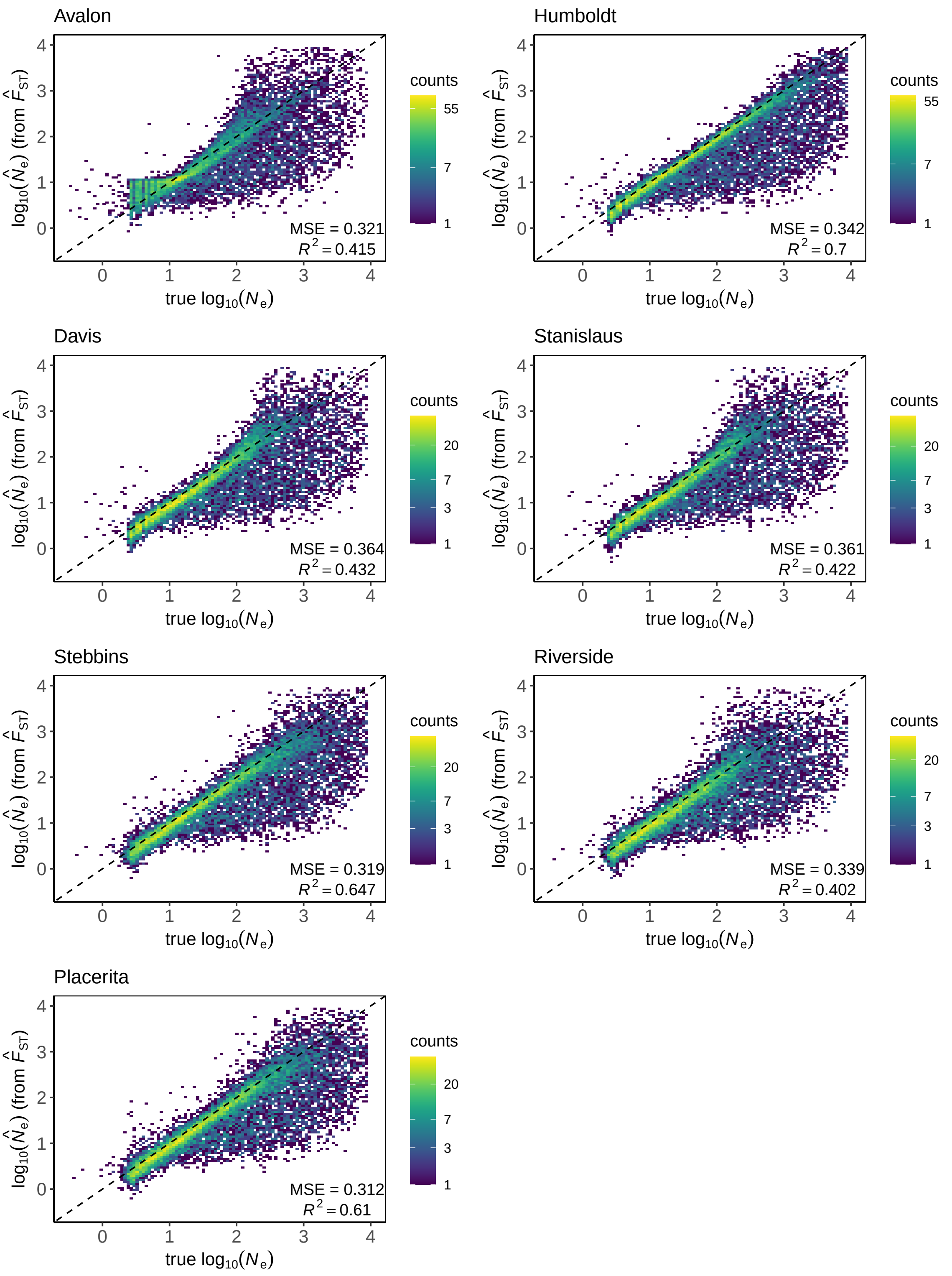


**Figure S26: $\hat{N}_e$ estimates from the temporal $F_{ST}$ to compare with the ABC-RF -based $\hat{N}_e$ estimates for pairs of temporal samples of *A. mellifera* feral populations.**

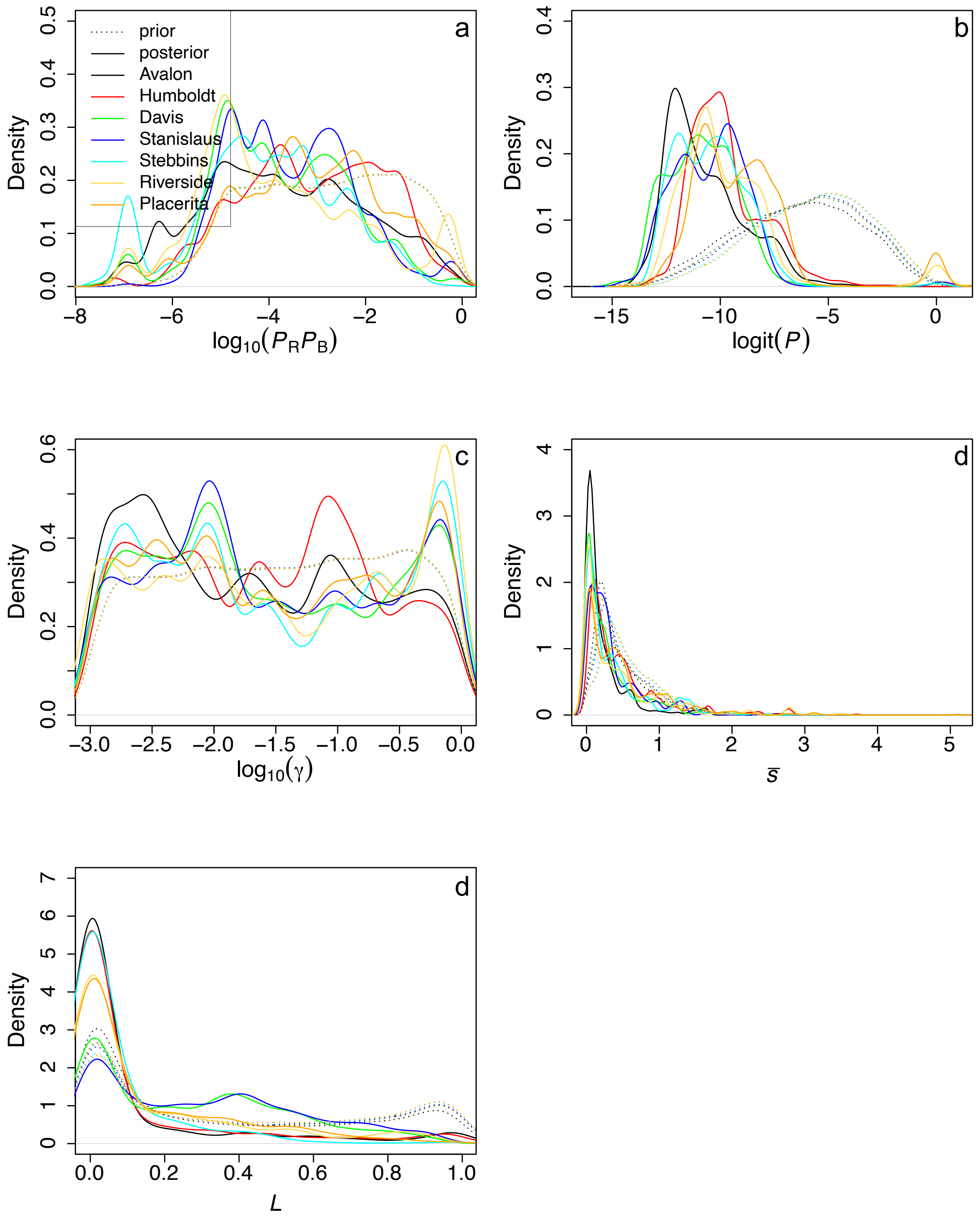


**Figure S27: Posterior distributions of model parameters and a latent variable informative about selection for all feral *A. mellifera* populations.** (a) probability of a beneficial mutation to arise $P_R P_B$; (b) number of mutations under strong selection $P$; (c) mean of the gamma distribution $\gamma$; (d) mean of selection coefficients of mutations under strong selection $\bar{s}$; and (e) mean substitution load $L$. Dashed and filled lines correspond to the prior and posterior distributions, respectively. See Table S3, Supplementary Results for mean estimates and 95% credibility intervals.

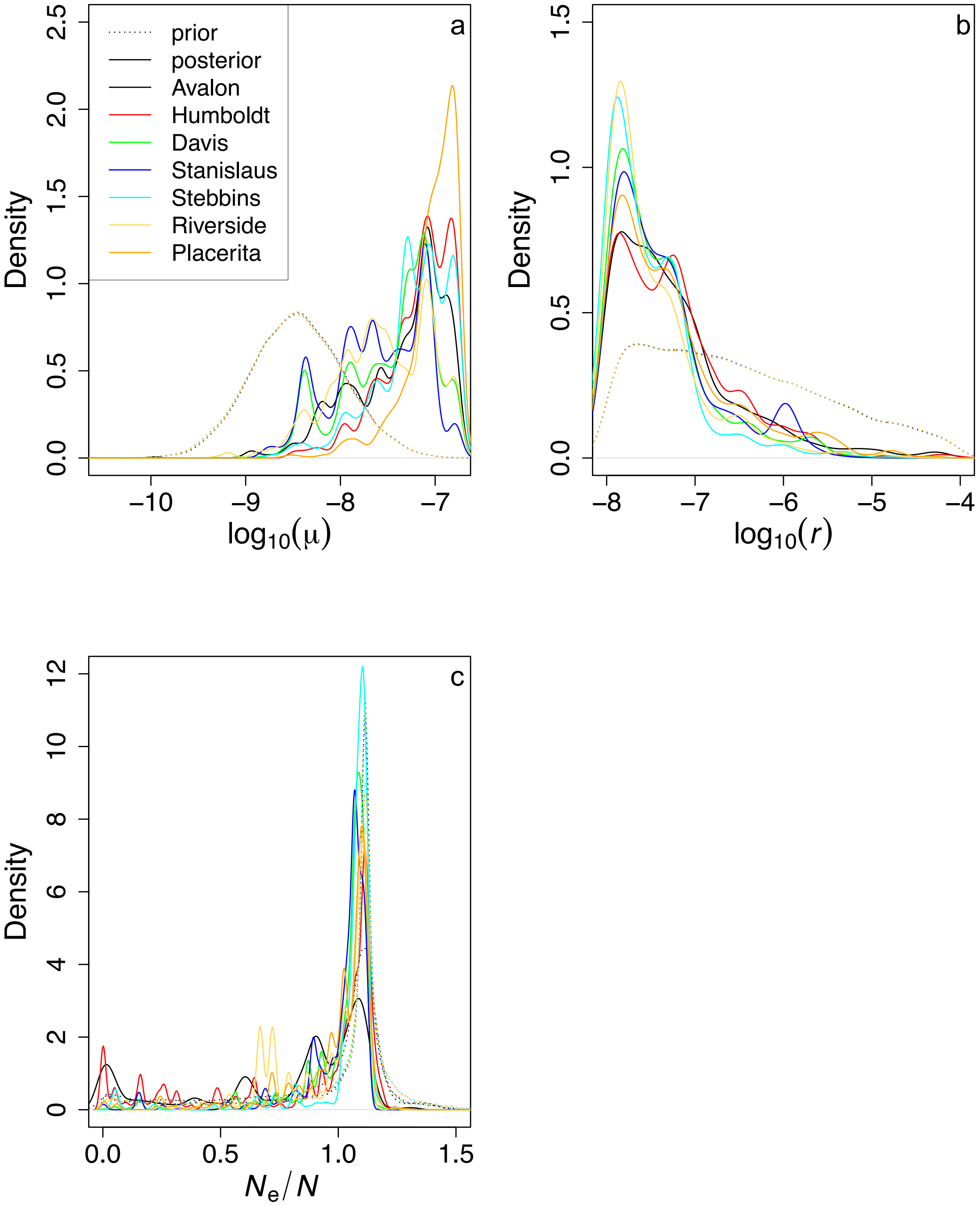


**Figure S28: Posterior distributions of model parameters and a latent variable informative about demography for all feral *A. mellifera* populations.** (a) mutation rate per generation $\mu$; (b) per base recombination rate per generation $r$; and (c) the ratio between the effective size and the population census size $N_e/N$. Dashed and filled lines correspond to the prior and posterior distributions, respectively. See Table S3, Supplementary Results for mean estimates and 95% credibility intervals.
